# Identifying Novel Targets of the Stringent Response in Plants and Cyanobacteria using chemoproteomics

**DOI:** 10.64898/2026.07.01.735797

**Authors:** Anna Karlsson, Rees Rillema, Emil Sporre, Elias Englund, Nikoleta Vogiatzi, Júlia Llavina Ramírez, Cenk O. Gurdap, Erdinc Sezgin, Fredrik Edfors, Cecilia Blikstad, Åsa Strand, Daniel C. Ducat, Elton P. Hudson

**Affiliations:** Science for Life Laboratory, School of Engineering Science in Chemistry, Biotechnology and Health, KTH Royal Institute of Technology, Stockholm, Sweden; Department of Energy-Michigan State University Plant Research Laboratories, Michigan State University, East Lansing, MI, United States; Department of Chemistry-Ångström Laboratory, Uppsala University, Uppsala, Sweden; Science for Life Laboratory, Department of Women’s and Children’s Health, Karolinska Institutet, Sweden; Umeå Plant Science Centre, Department of Plant Physiology, Umeå, Sweden

## Abstract

Survival in dynamic environments requires photosynthetic organisms to rapidly sense and respond to stress. The stringent response, mediated by the signaling molecule guanosine-3’,5’-bisdiphosphate (ppGpp), is crucial for acclimation to environmental changes such as darkness and nitrogen limitation. While it has been extensively characterized in heterotrophic bacteria such as *Escherichia coli*, the molecular mechanisms and regulatory targets of ppGpp in photosynthetic organisms remain less understood. Here, we report large-scale chemoproteomic identification of ppGpp-binding proteins across plant chloroplasts and cyanobacteria, revealing both conserved and novel targets compared to *E. coli*. In plants, we found that ppGpp regulates pyrimidine metabolism by inhibiting the chloroplastic enzyme aspartate transcarbamoylase (PyrB). In cyanobacteria, we found that ppGpp activates glucose-1-phosphate adenylyltransferase (GlgC) involved in glycogen synthesis, activates citrate synthase (GltA), and induces carboxysome aggregation. These findings expand the known ppGpp regulatory network in photosynthetic organisms and provide a foundation for understanding how ppGpp coordinates adaptation to nutrient and environmental stresses.

## Introduction

Secondary messenger molecules play essential roles in coordinating cellular responses to environmental changes. Guanosine-3’,5’-bisdiphosphate (ppGpp) mediates the stringent response, a global regulatory mechanism triggered during stress such as nutrient starvation. This response allows bacteria to conserve energy by downregulating growth-related processes, for example by inhibiting translation and transcription^1^. While the stringent response was first characterized in heterotrophic bacteria, it is conserved in photosynthetic cyanobacteria and plant chloroplasts. As an adaptation to photosynthetic life the stringent response is triggered by darkness, and further plays a role in regulating nitrogen metabolism^2^. Despite the central role of ppGpp, the molecular mechanism and regulatory targets of ppGpp in photosynthetic organisms are less understood compared to heterotrophic bacteria.

Cyanobacteria use the bifunctional SpoT enzyme to synthesize ppGpp. The cyanobacterial SpoT is activated in darkness and is essential for survival in day-night cycles^3^. In *Synechococcus elongatus*, accumulation of ppGpp was found to be induced after chemical inhibition of the photosynthetic electron chain by DCMU and DBMIB, but not by inhibition of isoleucyl-tRNA synthetase, disruption of the proton gradient, extreme pH, or high salt^4^. However, in *Anabaena* PCC 7210, ppGpp levels were reported to spike during serine hydroxamate-induced amino acid starvation^5^. Nitrogen limitation similarly causes ppGpp accumulation in several cyanobacteria species, and is essential for formation of nitrogen-fixing heterocysts in *Anabaena*^5–7^. Overexpression of ppGpp synthases in *Synechococcus* causes a chlorotic phenotype resembling nitrogen starvation^3,4,8^, and ppGpp is suggested to affect activity of the nitrogen regulatory PII protein by altering 2-oxoglutarate levels^9^.

In contrast to cyanobacteria, ppGpp synthesis in chloroplasts occurs via three classes of RSH proteins: RSH1, RSH2/3, and a calcium-binding RSH (CRSH)^2^. The plant RSH proteins differ structurally from the cyanobacterial enzyme, and phylogenetic analysis shows that RSH1 and CRSH cluster with the Deinococcus–Thermus lineage rather than with cyanobacteria, implying that plant RSH likely originated via lateral gene transfer rather than direct inheritance from the ancestral chloroplast^10^. RSH1 is homologous to the bacterial SpoT enzyme but functions exclusively as a ppGpp hydrolase that limits ppGpp levels under non-stress conditions. The paralogs RSH2 and RSH3 are bifunctional ppGpp synthethase/hydrolases with regions unique to photosynthetic eukaryotes. RSH2/RSH3 catalyse ppGpp synthesis during day-time stress and catalyse ppGpp hydrolysis during night. Recently, RSH3 activity was found to be directly regulated by the TOR nutrient and stress signaling pathway^11^. CRSH acts as a Ca²⁺-activated ppGpp synthase, rapidly accumulating ppGpp during dark transitions to adjust chloroplast gene expression to night conditions^12^. Interestingly, production of ppGpp is especially rapid in response to unexpected darkness compared to regular day-night cycles^8,13^. Similar to cyanobacteria, ppGpp is essential for acclimation to nitrogen limitation and contributes to the degradation of photosynthetic pigments^14,15^. In addition, plants have been found to produce ppGpp in response to a wide range of stresses, including wounding, heat shock, salinity, acidity, drought, UV irradiation, heavy metals, and pathogen attacks^13,16^.

Though the long-term effect of the stringent response has been studied through overexpression and knockdown studies^2^, no large-scale study has been made on direct binding targets of ppGpp in photosynthetic organisms. In heterotrophs like *E. coli*, *Staphylococcus aureus*, *Bacillus anthracis* and *Salmonella typhimurium*, screening approaches such as Differential Radial Capillary Action of Ligand Assay (DRaCALA) and affinity-capture have identified a wide range of ppGpp-binding proteins. These include GTPases involved in translation and ribosome biogenesis, metabolism of (p)ppGpp and derivatives, purine metabolism, and maturation of dehydrogenases^17–21^. From these studies, the importance of direct ppGpp-enzyme interactions in bacterial fitness has been explored. For example, ppGpp directly inhibits the nucleotide salvage enzyme Gsk, and mutating the enzyme Gsk to be ppGpp-insensitive disrupts nucleotide homeostasis and reduced starvation fitness in *E. coli*^22^.

To address the knowledge gap in photosynthetic organisms, and to compare differences between cyanobacterial and chloroplastidal stringent response, we applied a high-throughput chemoproteomic approach to identify ppGpp interaction partners.

Chemoproteomic methods detect potential ligand-protein interactions by measuring changes in protein stability or conformation upon binding^23^. These methods have been used to successfully identify drug and metabolite interactions in bacteria and plants^24–26^, and do not require prior knowledge of interaction partners or ligand design. We previously adapted Proteome Integral Solubility Alteration (PISA)^27^ for use in photosynthetic organisms^28^. PISA uses heat to partially denature proteins and measures changes in protein stability due to direct or indirect interaction with a ligand, allowing identification of interaction partners across the proteome via LC-MS.

Here we report large-scale screening of ppGpp-protein interactions across photosynthetic organisms using PISA: *Arabidopsis thaliana* chloroplasts, the cyanobacteria *Synechocystis sp.* PCC 6803 and *Synechococcus elongatus* PCC 7942. Our analysis identified multiple putative ppGpp targets and we characterized novel regulation of plant pyrimidine metabolism, cyanobacterial regulation of glycogen synthesis, and the CO₂-concentrating mechanism, as well as extend knowledge of ppGpp-mediated regulation of the citric acid cycle and nitrogen metabolism. This work provides new insights into the ppGpp regulatory network in photosynthetic organisms, offering a foundation for future research in stress adaptation.

## Results

### Screening of ppGpp-protein interactions using PISA

We performed PISA on cell extracts of *Synechocystis* sp. PCC 6803 *(Syn*6803) and *Synechococcus* sp. PCC 7942 (*Syn*7942), and extracts from *A. thaliana* chloroplasts with added ppGpp (Figure 1A). *E. coli* cell extracts were included to validate the method (Figure 1B), as ppGpp targets have been characterized in *E. coli*^1,18,19^, as well as to allow comparison with a heterotrophic strain. To limit metabolization of ppGpp in *Syn*6803, we used a previously generated mutant lacking the bifunctional ppGpp synthetase/hydrolase SpoT^29^.

**Figure 1.**
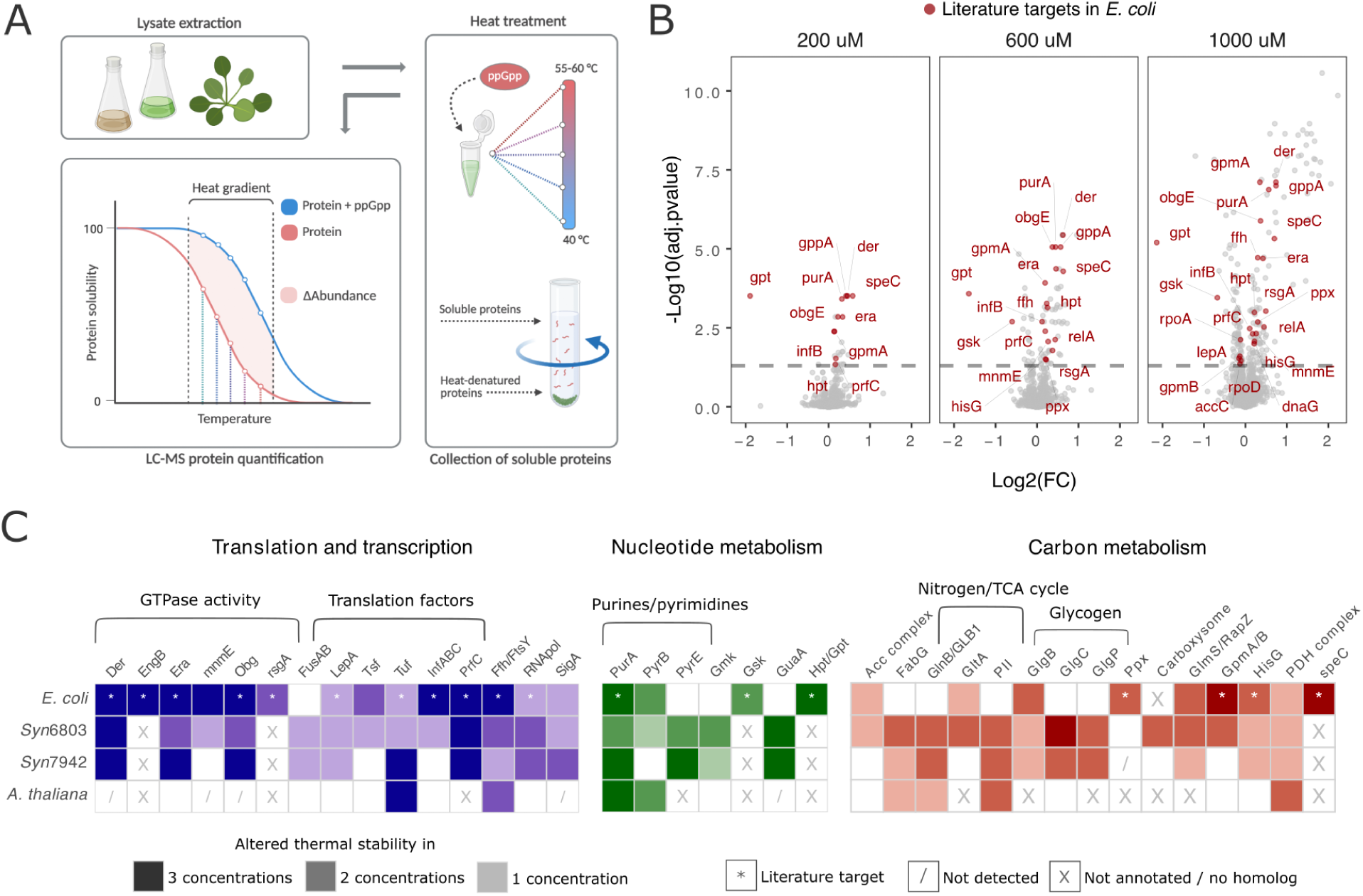
Screening of ppGpp-binding proteins using PISA. A) Overview of workflow. B) PISA method validation using ppGpp added to *E. coli* extracts. Previously reported ppGpp-binding targets in *E. coli* that showed an altered thermal stability in the presence of ppGpp are highlighted in red^1,18,19,32^. Treatments were with 200, 600 and 1000 μM ppGpp with technical quadruplicates, compared against a control without ppGpp added. C) Proteins with significantly (adjusted p.value ≥ 0.05) affected thermal stability in PISA for the tested organisms. The tested concentrations of ppGpp were 200, 600 and 1000 μM with technical quadruplicates compared to a control without ppGpp.

To perform the PISA experiment, cultures of *E. coli*, *Syn*7942 and *Syn*6803 were harvested during the exponential phase and lysed. For *A. thaliana*, chloroplasts were extracted from leaves during the end of subjective night. Before addition of ppGpp, endogenous metabolites were removed from the crude proteome by desalting via size exclusion columns. A minimum ppGpp concentration of 200 μM concentration was used based on an estimate of maximal intracellular ppGpp levels in dark-treated cyanobacteria^3,9^, while 1 mM corresponds to the intracellular concentration of ppGpp during the full stringent response in *E. coli*^30^. Samples were then incubated at a heat gradient to induce protein denaturation, and soluble, non-denatured proteins were recovered by ultracentrifugation. *E. coli* and cyanobacteria cell extracts were incubated at 40-60 °C, and chloroplast extracts at 40-55 °C, temperature ranges selected based on reported proteome stability^31^. LC-MS was used to quantify changes of soluble protein, corresponding to a shift in melting temperature with ppGpp present in the sample compared to the control without ppGpp. Proteins that present a shift in thermal stability (adjusted p-value ≥ 0.05) are potential interaction partners with ppGpp, and proteins showing a consistent decrease or increase in stability across multiple ppGpp concentrations are more likely to represent true positives^27,28^. The LC-MS measurements had a total proteome coverage of 2143, 1594, 1868 and 1286 proteins for *Syn*6803, *Syn*7942, *E. coli* and *A. thaliana* chloroplasts, respectively. At 600 μM ppGpp, we detected 183 (*Syn*6803), 63 (*Syn*7942), 78 (*E. coli*) and 52 (*A. thaliana*) proteins with shifted thermal stability (adjusted p-value < 0.05). Notably, the Uniprot reference proteome for *Syn*7942 contains 850 fewer entries and fewer protein annotations than the *Syn*6803 reference proteome. Complete lists of proteins with an altered melting temperature for each ppGpp concentration are available in Supplementary File 1.

To validate the ability of PISA to identify true interactions, we compared affected *E. coli* proteins with previously characterized ppGpp targets identified using DRaCALA and capture-compound mass spectrometry (Figure 1B)^1,18,19,32^. We detected many of the known ppGpp-binding proteins, including GTPases involved in translation (Der, Era, Obg, RsgA), translation factors (LepA, Tuf, Inf B, prfC), RNA polymerase subunits, nucleotide metabolism (PurA, Gsk, Hpt, Gpt) and ppGpp-metabolism (gppA, relA)^1,30^. Multiple ribosomal subunits and tRNA ligases have affected thermal stability by ppGpp, likely due to nonspecific, indirect binding of ppGpp via GTPases and translation factors. In *E. coli*, ribosomal subunits accounted for 25% of the total proteins with altered thermal stability. However, some ribosomal subunits may play important roles in ppGpp-dependent regulation. For example, the 30S ribosomal protein S2 (RpsB), which showed altered stability in our screen in both *Syn*6803 and *Syn*7942, has also been reported to mutate in ppGpp-deficient Syn7942 strains^33^.

Cyanobacteria and *E. coli* shared several common targets in our screening, including GTPases involved in translation (Der, Era, Obg) and translation factors (PrfC, InfA/B/C, LepA, FusA/B, EF-tu/tuf) (Figure 1C). This indicates that regulation of translation via GTPases is a conserved function of ppGpp, which agrees with previous studies across gram-positive and gram-negative bacteria^1,30^. Fewer translation-related proteins were identified in the *A. thaliana* screen. Several bacterial GTPases lack clear annotated homologs in plants, and some of the chloroplastidial bacteria-like GTPases were not detected in our dataset. However, the elongation factor tu (EF-tu/tufA) showed altered stability in our dataset across both bacteria and plants. Moreover, plant GTPases involved in post-translational modification and protein transport were affected, such as BPG2, SAR1A/B, and ARF1/2. Notably, ppGpp affected the signal recognition particle (SRP) complex in all organisms, which is important for membrane protein insertions and a conserved target for the stringent response in *B. subtilis* and *E. coli*^32^. In plants, SRP is essential for insertion of light-harvesting chlorophyll-binding proteins into the thylakoid membrane^34^. Overall, GTPases detected in our datasets are promising candidates for ppGpp interaction, as they play key regulatory roles in cellular processes and are known targets of ppGpp in other bacteria.

### ppGpp effects on RNA polymerase

RNA polymerase subunits and the primary sigma factor A were affected by ppGpp in *E. coli* and in both tested cyanobacteria. In *E. coli*, ppGpp is known to directly interact with RNA polymerase at a site in between the β′ (rpoC) and ω (rpoZ) subunits, and at a second site where ppGpp binds together with the transcription factor DksA^35^. However, the importance of ppGpp binding to RNA polymerase is unclear, as removal of the ppGpp binding site on RNA polymerase had little effect on transcription profile^19^. Cyanobacteria RNA polymerase is structurally different from the *E. coli* enzyme, and contains β′ and ω subunits, but lacks a homolog for DksA, and direct binding of ppGpp has not been studied. In *A. thaliana*, we did not detect changes in the stability of bacteria-like plastid-encoded RNA-polymerase (PEP) or non-bacterial PEP-associated proteins in chloroplasts. Similar to cyanobacteria, the PEP complex does not have the ppGpp binding sites present in *E. coli* and lacks both DksA and the ω subunit. However, previous studies have shown ppGpp inhibition of the PEP complex by likely binding the β′ subunit in tobacco and red algae, though at weaker sensitivity than *E. coli* RNA polymerase^36,37^.

### ppGpp inhibits pyrimidine metabolism in plants

One way the stringent response reduces growth is by limiting production of new nucleotides. Adenylosuccinate synthetase (purA) is a key step in purine biosynthesis and was previously shown to be a ppGpp target in *E. coli*^19^. PurA was a conserved target of the stringent response based on our screening across all organisms (Figure 1C). We also detect the purine metabolism enzyme GuaA as affected by ppGpp, consistent with a recent suppressor genetic screen in *Syn*7942^33^. In addition to purine metabolism, we found that pyrimidine metabolism is also regulated by ppGpp. Aspartate carbamoyltransferase (PyrB) was altered by ppGpp in plant chloroplasts. PyrB catalyzes the first step of *de novo* pyrimidine synthesis in the chloroplast and is growth-limiting in *A. thaliana*^38^. Later steps in the pyrimidine synthesis pathway are carried out in the plant cell cytosol and are therefore unlikely to be regulated by ppGpp.

To confirm ppGpp as a regulator of *A. thaliana* PyrB activity, we recombinantly expressed and purified the protein. Using Nano Differential Scanning Fluorimetry (NanoDSF), the melting temperature of PyrB was reduced by 17 °C upon addition of ppGpp, indicating a direct interaction (Figure 2A). Further investigation by kinetic assays revealed inhibition of PyrB in the presence of 200 μM ppGpp (Figure 2B). We did not observe an effect on catalytic activity by GDP, the metabolic precursor to ppGpp, indicating the effect is specific to ppGpp (Supplementary Figure 1A).

**Figure 2.**
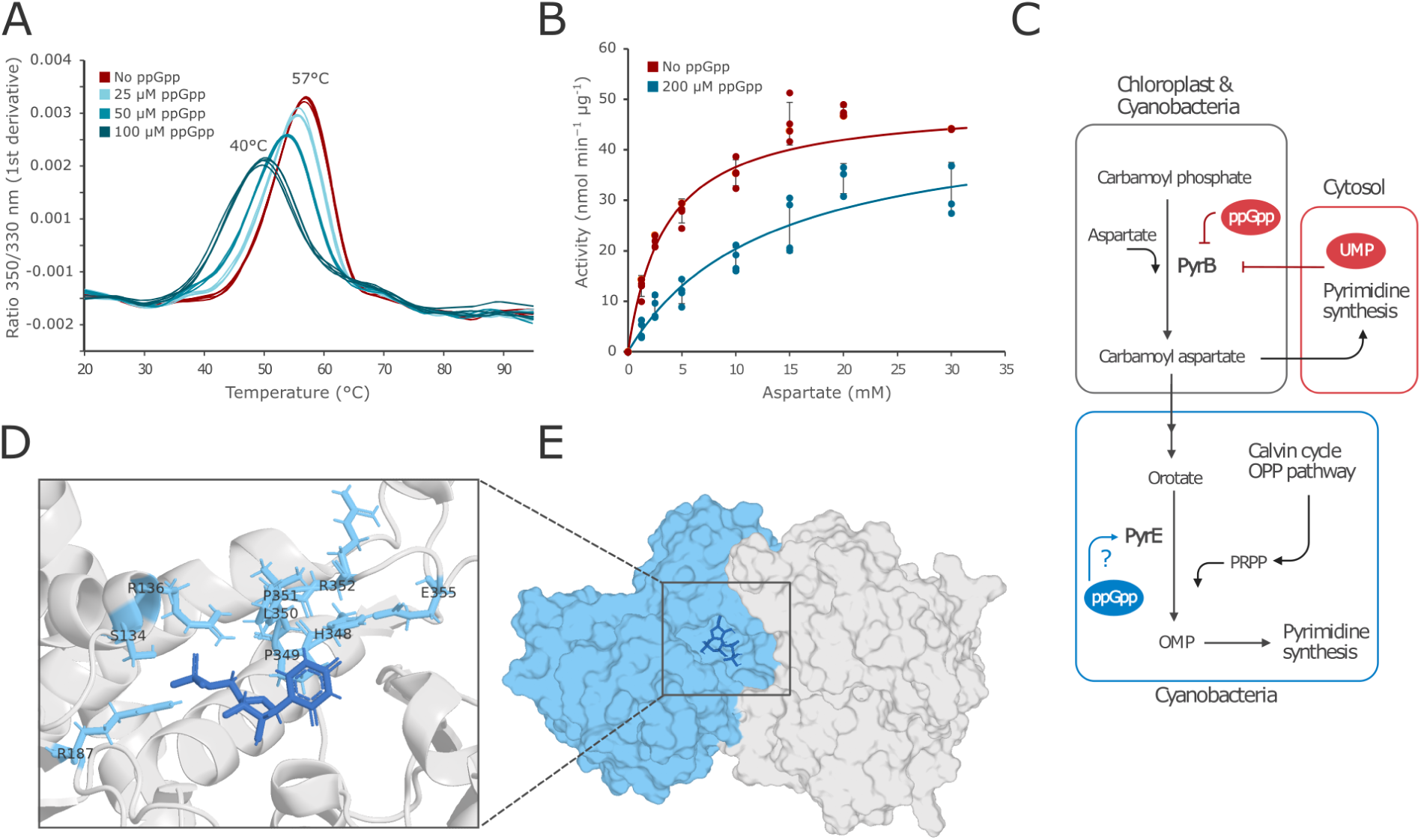
Validation of ppGpp interaction with recombinantly expressed *Arabidopsis thaliana* PyrB. A) Shift in thermal stability in the presence of ppGpp in quadruplicate measurements. B) Kinetic assay showing inhibition by ppGpp. The Michaelis–Menten equation was fitted to the data using non-linear regression. Lines represent the mean and standard deviation (SD) of four replicates. C) Overview of pyrimidine metabolism and regulation by ppGpp. In plants, later steps in pyrimidine synthesis are carried out in the cytosol. D) Residues in the active site (within 5 Å) affected by limited proteolysis (light blue) with 200 μM ppGpp (adjusted p-value <0.05), shown in an monomer crystal structure with UMP bound (PDB: 6YPO). UMP is shown in dark blue. E) DynamicBind docking of ppGpp using the dimeric UMP-bound crystal structure (PDB: 6YPO, UMP molecule excluded) with ppGpp (dark blue) docked in the active site.

The binding site of ppGpp to the *A. thaliana* PyrB was investigated by limited proteolysis small-molecule mapping on cell lysates spiked with purified PyrB protein^25,39^. Using this method, we found residues in the carbamoyl phosphate binding pocket to be affected by ppGpp, suggesting that this region may form a ppGpp binding site similar to the binding position of the plant-specific feedback inhibitor UMP^38^ (Figure 2C). These results were consistent with a docking analysis of ppGpp to the crystal structure of UMP-bound PyrB (Figure 2D). Predictions generated during docking were located in the same binding pocket, regardless of using the apoform (PDB: 6YY1) or UMP-bound crystal structure of PyrB (PDB: 6YPO).

In *Syn*6803, PyrB only had an altered stability at the highest tested ppGpp concentration and not at all in *Syn*7942. Instead, orotate phosphoribosyltransferase (PyrE) was strongly affected in the PISA screen in both *Syn*6803 and *Syn*7942. PyrE catalyses an intermediate step in pyrimidine synthesis using 5-phosphoribosyl-1-pyrophosphate (PRPP) as a substrate and links ribose metabolism to pyrimidine metabolism. This could be an alternative regulatory point in cyanobacteria. To validate that ppGpp directly affects PyrE, we cloned and purified the *Syn*6803 PyrE and tested ppGpp interaction using NanoDSF. The PyrE melting point was increased by 5 °C in the presence of ppGpp, indicating binding (Supplementary Figure 1B). Other enzymes with PRPP binding pockets, such as XPRT, PurR, PurF, are known to bind ppGpp competitively, and makes PyrE a likely target for ppGpp inhibition^19,40,41^.

### ppGpp activates glycogen synthesis in cyanobacteria

Enzymes involved in glycogen metabolism were affected by ppGpp in the bacterial species in our PISA assay but not in chloroplast. Plants and cyanobacteria share several enzymes for glycogen and starch synthesis (Figure 3A)^42^, and both starch and glycogen accumulate during nitrogen starvation. However, unlike cyanobacteria, which accumulate glycogen in ppGpp overexpression mutants^9^, ppGpp overproduction in plants does not increase starch levels, and blocks starch accumulation during nitrogen starvation^43^. We therefore sought to validate the effect of ppGpp on glycogen and starch metabolism enzymes.

**Figure 3.**
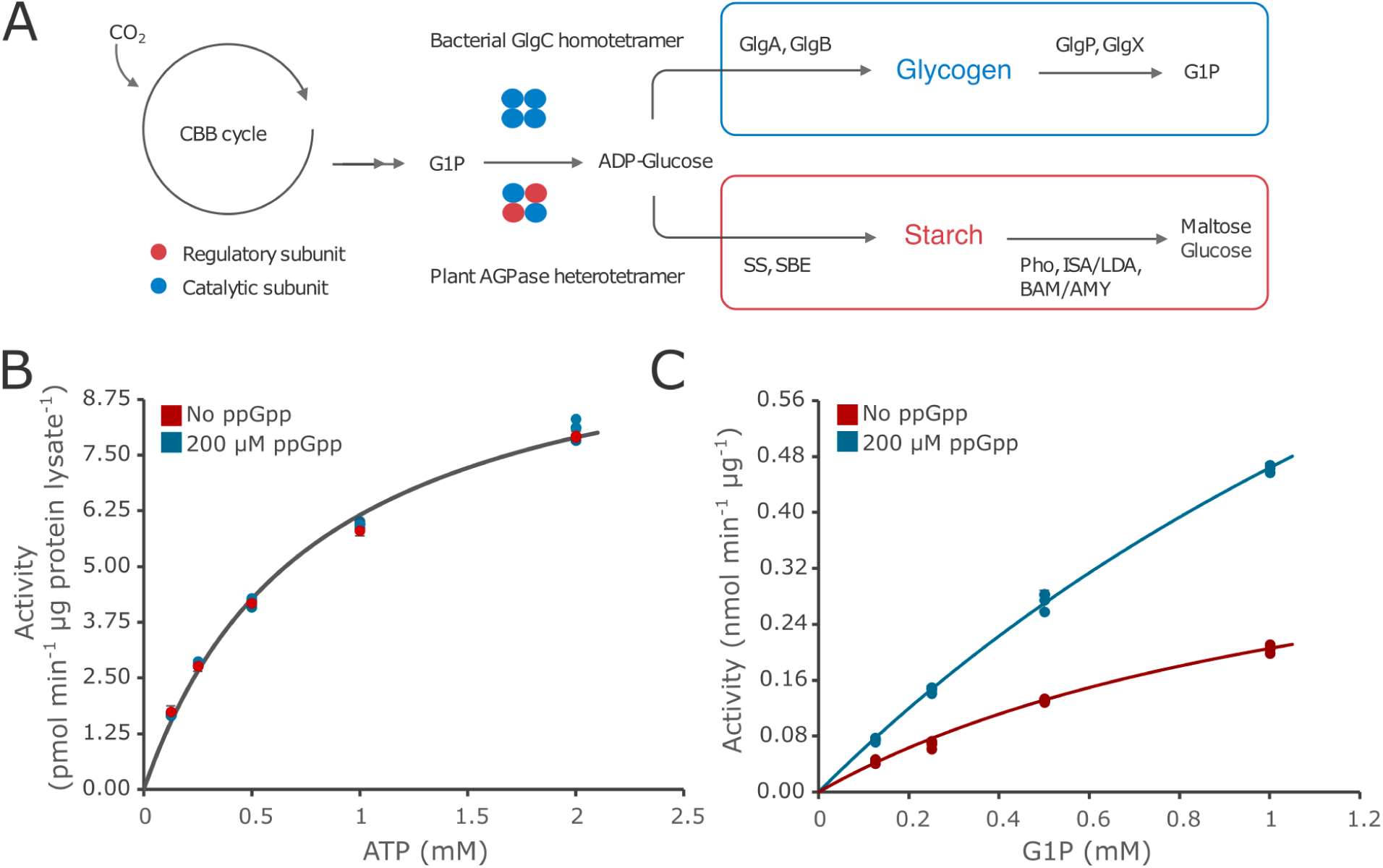
Interaction with ppGpp in glycogen synthesis. A) Overview of starch and glycogen metabolism in cyanobacteria and plants. B) Assay of glucose-1-phosphate adenylyltransferase activity, measuring formation of ADP-glucose from ATP and glucose-1-phosphate in *A. thaliana* leaf lysates, and C) using recombinant purified *Syn*6803 GlgC. The Michaelis–Menten equation was fitted to the data using non-linear regression. Lines represent the mean ± SD of three replicates. Abbreviations: G1P, glucose-1-phosphate; SS, starch synthases; SBE, starch branching enzymes; Pho, starch phosphorylase; ISA/LDA, debranching enzymes; BAM/AMY, amylases.

To assess starch synthesis rates in leaf extracts, we measured ADP-glucose formation (the precursor of starch) catalyzed by glucose-1-phosphate adenylyltransferase, using UV-HPLC (Supplementary Figure 2A). Addition of 200 µM ppGpp to leaf extracts had no detectable effect on ADP-glucose synthesis (Figure 3B). It was not possible to assess glycogen accumulation in cyanobacteria extracts, as glucose-1-phosphate adenylyltransferase activity was too low to measure. We therefore expressed *Syn*6803 GlgC recombinantly and measured its activity, and observed an activation of the enzyme in the presence of 200 µM ppGpp (Figure 3C). These results are in agreement with the PISA data, cyanobacteria glycogen synthesis is affected by ppGpp, while plant starch synthesis was not.

Lastly, we observed that Amy3, a redox-regulated α-amylase involved in stress-induced starch degradation^44^, displayed altered thermal stability in chloroplasts. However, *in vitro* starch degradation assay with recombinant Amy3 revealed no direct effect of ppGpp (Supplementary Figure 2B).

### Effects of ppGpp on nitrogen assimilation and the TCA cycle

The stringent response is needed for nitrogen starvation acclimation in plants and *E. coli*, and ppGpp overexpressing cyanobacteria typically have a chlorotic phenotype^3,4,15,45,46^. Across all photosynthetic organisms, the regulatory PII protein (GlnB/GLB1) was affected by ppGpp. The PII protein regulates nitrogen metabolism based on ATP/ADP ratio, and 2-oxoglutate levels^47^. In *E. coli* extracts, the related PII-like protein GlnK was affected, but not GlnB. To test if ppGpp could directly interact with the PII protein, we purified the *Syn*6803 GlnB recombinantly and measured binding affinity of ppGpp via Isothermal Titration Calorimetry (ITC). No interaction was observed (Supplementary Figure 3), and we concluded that any potential effect on the nitrogen regulatory protein PII (GlnB) is indirect.

In cyanobacteria, it is proposed that ppGpp-regulation of PII is mediated through the reduction in 2-oxoglutarate, potentially through direct kinetic inhibition of the TCA cycle enzyme aconitase in *Syn*7942^9^. We saw weak evidence of direct aconitase-ppGpp interaction at 1 mM ppGpp in *Syn*6803, and not in *Syn*7942 (adjusted p-value of 0.1 at 1 mM ppGpp). Instead, we found the enzyme citrate synthase (GltA), the enzyme preceding aconitase in the TCA cycle, to have increased thermal stability upon ppGpp addition in *Syn*6803. We did not observe a change in *Syn*7942 GltA (Figure 1C), consistent with cell lysate assays previously reported^9^.

We investigated *Syn*6803 GltA by recombinant expression in *E. coli* and verified an effect on the thermal stability on purified protein in the presence of ppGpp via NanoDSF (Figure 4B). In the presence of 5 mM MgCl_2_, a known activator of *Syn*6803 GltA^48^, a small but reproducible shift in melting temperature was observed (1 °C). Mass photometry analysis of the enzyme’s oligomerization state also revealed a shift from hexamers to dimers in the presence of ppGpp (Figure 4C). To investigate if ppGpp affects the catalytic properties of GltA, we did an *in vitro* enzymatic assay which revealed a modest increase in activity in presence of 200 μM ppGpp (Figure 4D). The increase in enzymatic activity was present both with and without MgCl_2_ (Supplementary Figure 4A). Since *Syn*6803 GltA is known to be affected by various salts^48^, we controlled for the effect of lithium ions present in the commercial ppGpp salt. However, addition of LiCl had no effect on the activity of GltA (Supplementary Figure 4B).

**Figure 4.**
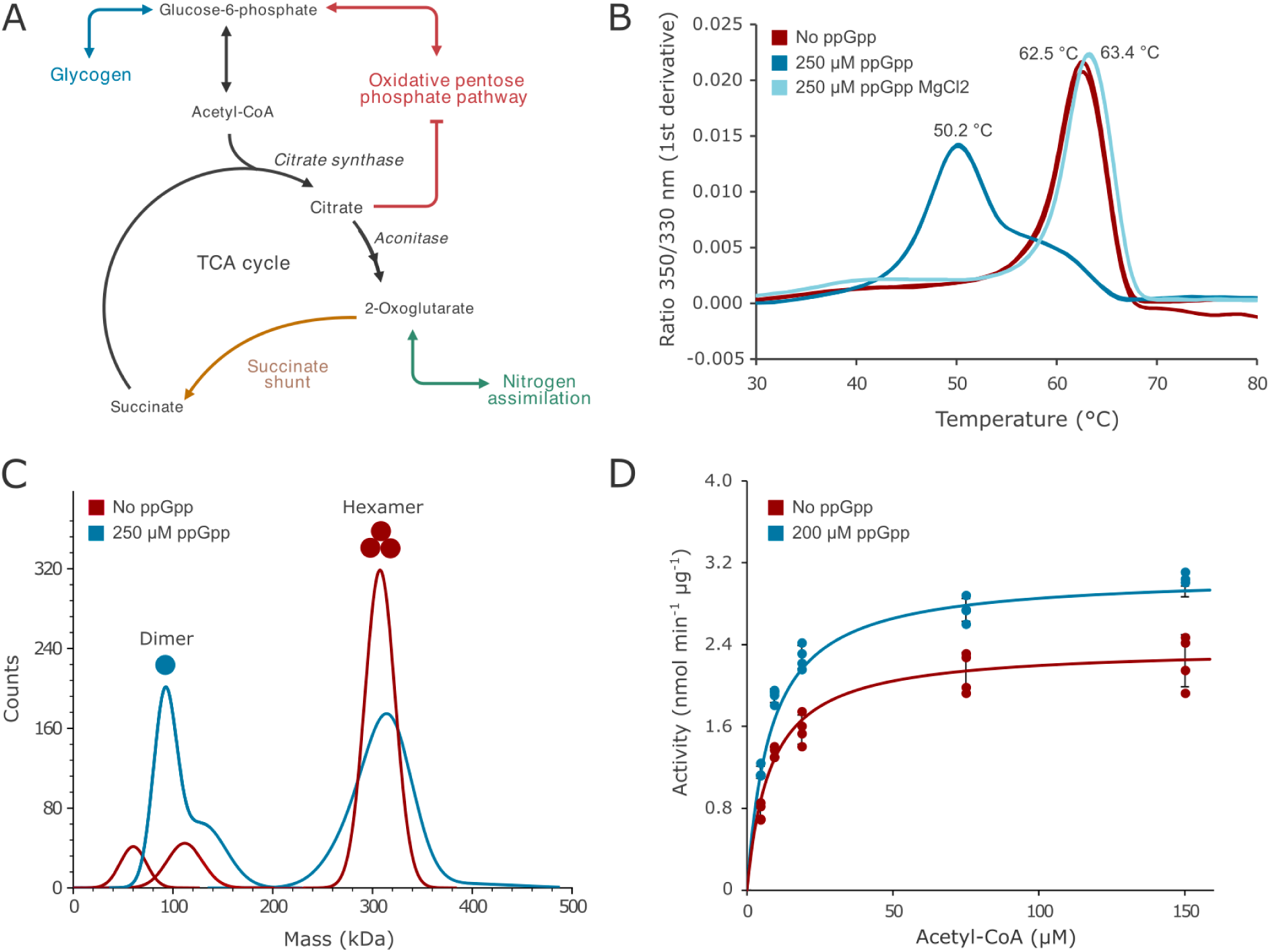
A) Overview of citrate synthase and related metabolism in cyanobacteria. B) Shift in thermal stability of *Syn*6803 citrate synthase (GltA) in the presence of ppGpp using triplicate measurements. C) Mass photometry of GltA showing a shift from hexamers to dimers in the presence of ppGpp with 5 mM MgCl_2_. D) Activation of GltA by ppGpp in the presence of 5 mM MgCl_2_. The Michaelis-Menten equation is fitted to the data using non-linear regression. Lines represent the mean and SD of four replicates.

### The stringent response induces carboxysome aggregation *in vivo*

PISA screening in *Syn*6803 revealed multiple carboxysomal subunits with altered thermal stability; The carboxysome shell proteins CcmL (affected at 2 concentrations of ppGpp), CcmP, CcmK1, CcmK2, CcmK3, CcmO (1 concentration), and the scaffolding protein CcmM (1 concentration). CcmL, showing the strongest response to ppGpp, is a pentameric protein that caps the corner of the carboxysome.

To investigate ppGpp interaction with *Syn*6803 carboxysome subunits, we measured the shift in thermal stability on purified *Syn*6803 carboxysome proteins; CcmL, CcmP, CcmK1, CcmK2, and CcmK4 (forming the carboxysome shell), CcmN (scaffolding protein), and carbonic anhydrase CcaA. Of the tested carboxysome proteins, only CcmL and CcaA produced a reliable melting curve. Addition of ppGpp altered CcmL melting temperature from 78 °C to 90 °C (Figure 5A), and changed the melting curve profile, which suggests a structural change induced by ppGpp that alters exposure of tryptophan residues measured by NanoDSF. Stability of recombinant CcaA was not affected by ppGpp (Supplementary Figure 5). In contrast, PISA analysis in *Syn*7942 extracts did not show detectable stability changes in any of the carboxysome proteins upon addition of ppGpp. This was confirmed by NanoDSF of purified *Syn*7942 CcmL (Supplementary Figure 5).

**Figure 5.**
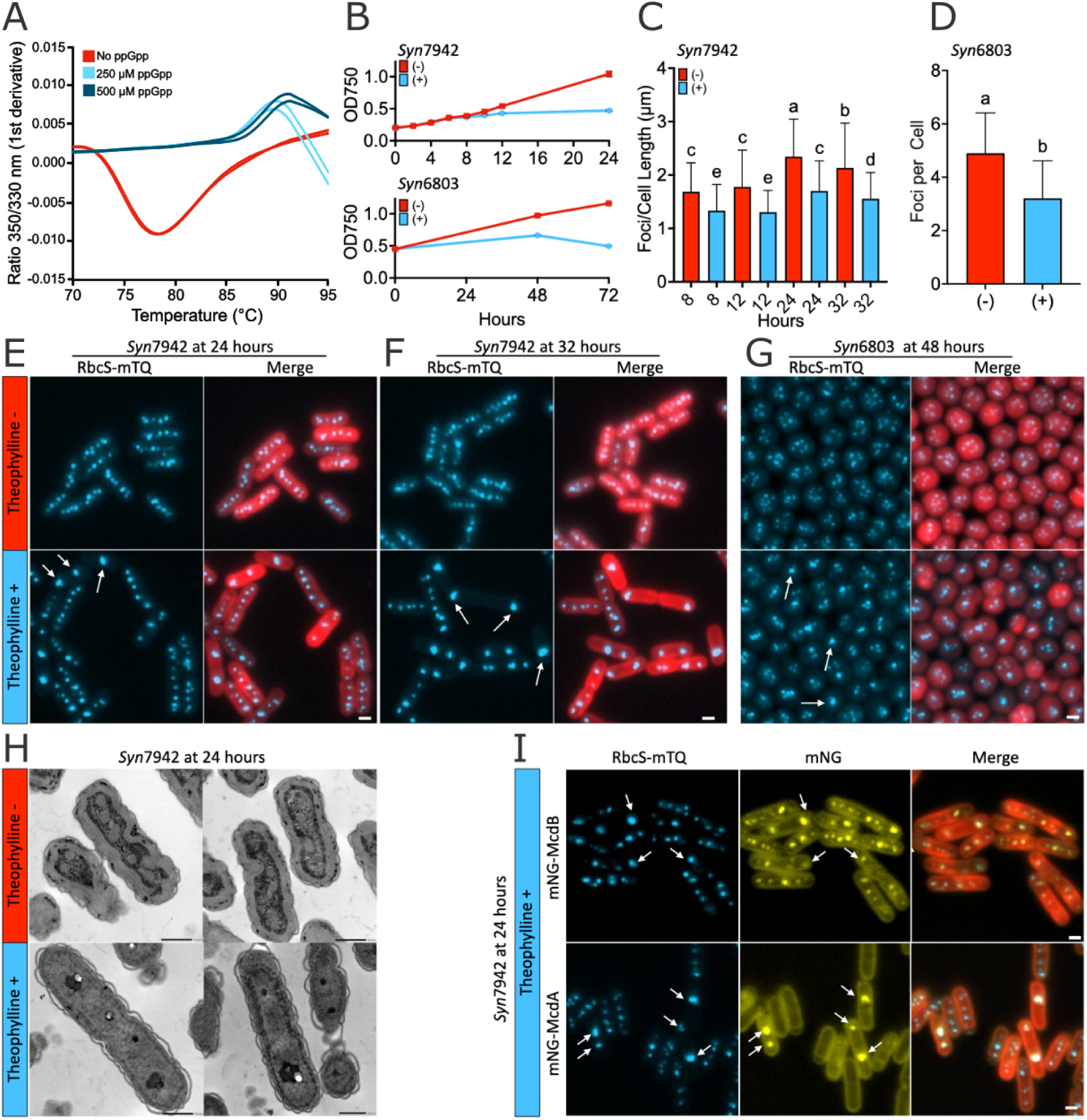
The effect of ppGpp on carboxysomes. A) ppGpp causes a shift in thermal stability of purified carboxysome protein ccmL in *Syn*6803. Measurements were made in duplicates. B) Growth curve of *Syn*7942 (top) and *Syn*6803 (bottom) expressing RelQ (YjbM) under a theophylline-inducible riboswitch element; with (+) and without (−) theophylline addition (1 mM). Symbols represent mean and SD of three biological replicates. C,D) Number of carboxysome foci (RbcS-mTurquioise2) per cell length of *Syn*7942 C) and per cell of *Syn*6803 in a time series following induction of YjbM with (+) and without (−) theophylline addition (1 mM). C,D) Bars represent mean and SD, n>472 cells. E and F) Fluorescence microscopy imaging of carboxysome foci using RbcS-mTurquoise2 at 24, and 32 hours following YjbM induction in *Syn*7942. G) Fluorescence microscopy imaging of carboxysome foci using RbcS-mTurquoise2 at 48h hours following YjbM induction in *Syn*6803. H) TEM imaging of RbcS-mTurquoise2 YjbM *Syn*7942 24 hours with (+) or without (−) induction of YjbM. I) mNeonGreen-McdB and mNeonGreen-McdA signal overlaps with the fluorescence signal of RbcS-mTurquoise2. Cells with severely mispositioned carboxysomes after 24 hours with induction of YjbM (arrowheads).

To investigate if ppGpp affects carboxysomes *in vivo* we utilized *Syn*7942 and *Syn*6803 strains expressing fluorescently-tagged Rubisco (RbcS-mTurquoise2), allowing visualization of carboxysomes due to Rubisco’s encapsulation within the organelle^49–51^. In these strains, we expressed the *B. subtilis* small ppGpp synthase RelQ (YjbM) under an theophylline-sensitive promoter. Upon induction of ppGpp synthesis and in line with previous studies in cyanobacteria, our ppGpp overproduction *Syn*7942 and *Syn*6803 strain showed growth arrest and degradation of photosynthetic pigments, with 50% lower Chlorophyll *a* and carotenoid content after 24 hours (Figure 5B and Supplementary Figure 6). Fluorescence microscopy revealed that the number of detectable carboxysome foci decreased over time in samples induced to accumulate ppGpp (Figure 5C and D). Yet this reduction in pigment, growth, and carboxysomes was not associated with cell death (Supplementary Figure 6). Rather, carboxysome foci appeared to aggregate, with cells displaying an increasing number of large foci and decreased regularity of carboxysome spacing over time (Figure 5E-G). In previous reports of irregular rubisco foci, this phenotype is associated with carboxysome aggregation rather than a merging of carboxysome into a single conglomerate. Such mispositioning can also be associated with a reduction in carbon fixation via Rubisco in *Syn*7942^49–51^. We confirmed that these foci in *Syn*7942 represent distinct carboxysomes within an aggregate by transmission electron microscopy (TEM; Figure 5H). Interestingly, carboxysomes in the ppGpp-overexpressing strain also consistently displayed increased density of staining with the TEM dye, possibly suggesting alterations in carboxysome porosity or other physical features.

Cyanobacteria change ppGpp concentration within the cell in response to circadian rhythms, particularly accumulating ppGpp in the dark. This signals the cells to wait for anticipated light by arresting cell growth and inhibiting protein synthesis^3,4^. Prolonged dark has also been shown to inhibit the McdAB system, which is responsible for equidistantly spacing carboxysome along the axis of cell^52^. To determine if carboxysome aggregation caused by ppGpp is due to a generalized downregulation/inactivation of McdAB, we tagged McdB and McdA with a fluorescent reporter^53^. We observe that McdB is still strongly localized to the vicinity of carboxysome aggregates, suggesting that any structural changes to the carboxysome following ppGpp exposure does not lead to a dissociation of positional machinery or otherwise downregulate key carboxysome positional system components (Figure 5I). McdA is the oscillatory component of the position system, yet we see that McdA localizes to carboxysomes with ppGpp exposure (Figure 5I). Together, these results indicate that ppGpp modulates both the stability of specific carboxysome proteins and the spatial organization of carboxysomes *in vivo*, potentially representing a regulatory mechanism linking cellular stress responses to CO₂ fixation machinery. However, we did not observe an effect on the McdAB positional system in the PISA assay.

## Discussion

ppGpp is a key global regulator of metabolism and deeper understanding of ppGpp signaling could help guide biotechnological and agricultural applications. Modulating ppGpp levels can enhance substrate uptake rates and increase cellular robustness in large-scale industrial cultivations in *E. coli*, particularly under fluctuating nutrient conditions^54^. In plants, engineering ppGpp levels may improve tolerance to nutrient limitation and high light stress^43,55^. Targeting the stringent response, such as with synthetic ppGpp analogs, also offers a promising strategy for antimicrobial development^56^.

PISA is a powerful method for studying protein–metabolite interactions that does not require ligand design, as in pull-down assays, or prior knowledge of potential interactions. In this study, we found new ppGpp interactors and successfully detected several known ppGpp interactors previously characterized in *E. coli* (Figure 1C). However, some reported targets (HflX, BipA, Apt, HypB) were not identified as interactors in our screen, and others (XpT, YgdH, PpnN, SpeF) were not detected by LC-MS^2,30^. Because PISA relies on detecting ligand-induced changes in protein thermal stability, small shifts or melting points occurring outside the heat gradient used here (40–55/60 °C) can lead to false negatives. However, expanding the temperature span can reduce the signal-to-noise ratio, creating a trade-off in assay design^57^. PISA can also detect indirect interactions through co-melting, which is advantageous for probing large complexes and interaction networks but may introduce false positives that complicate downstream validation. Additionally, by incorporating a metabolite gradient, PISA can give insight into high- and low-affinity interactions, which is of interest for molecules such as ppGpp where the metabolic response depends on concentration^30^. Overall, overlapping screening techniques in combination with targeted studies are required for a full picture of the stringent response, but this study addresses an important gap in current knowledge and highlights proteins for further studies.

A novel finding of this study is post-translational regulation of pyrimidine metabolism by ppGpp. Based on our screening, we saw direct regulation of the plant enzyme PyrB that catalyses the initial step in pyrimidine metabolism in chloroplasts. Downregulation of PyrB in *Arabidopsis* stunts growth, and decreases photosynthetic efficiency^38^, similar to phenotypes seen in ppGpp overexpression lines^2^. The inhibition by ppGpp shows similarity with previously characterized UMP inhibition in plants^38^. Notably, UMP shifts the plant enzyme into a conformation that reduces catalytic activity drastically at low aspartate levels (<1 mM)^38^, below the substrate levels tested in this study.

PyrB structure and regulation vary considerably across organisms, raising the question of whether ppGpp inhibition is conserved. In our screen, ppGpp affected the stability of the *Syn*6803 and *E. coli* homologs of PyrB, but current knowledge of bacterial PyrB differs substantially from that of the plant enzyme. The *E. coli* PyrB is composed of a catalytic PyrB subunit and a regulatory PyrI subunit regulated by CTP and ATP. In contrast, PyrB in *B. subtilis* and many other bacteria is insensitive to direct nucleotide regulation, as they lack PyrI^58^. Similar to the *B. subtilis* enzyme, cyanobacteria lack the PyrI regulatory subunit, and while plant PyrB is similar to the cyanobacterial enzyme, the regulatory loop that induces an inhibited conformation in the presence of UMP is thought to be unique to plants^38^. To our knowledge no characterization of the cyanobacterial enzyme has been reported.

This study also shows activation of cyanobacterial GlgC by ppGpp, which catalyzes a rate-limiting step in glycogen synthesis. Accumulation of glycogen under high ppGpp levels has been reported previously in cyanobacteria and *E. coli*^9,59^. Consistent with our PISA screen, we observe activation of the cyanobacterial enzyme but not its plant homolog. The direct stimulation of GlgC activity by ppGpp provides a further mechanistic link connecting the hyperaccumulation of glycogen in cyanobacteria that is observed under some environmental stresses, such as nitrogen depletion. The plant enzyme likely evolved from the homotetrameric cyanobacterial GlgC and retains regulation by 3-phosphoglycerate and inorganic phosphate, but has since become a heterotetramer with a redox-regulated subunit^42,60^. Although glycogen and starch synthesis share core enzymes, these regulatory differences may reflect distinct stress responses, as ppGpp overproduction does not induce starch accumulation in plants^43^. Unlike the plant and cyanobacterial enzyme, the *E. coli* enzyme is regulated by fructose-1,6-bisphosphate and AMP. The *E. coli* GlgC has been reported to be inhibited by ppGpp at low levels of the substrate glucose-1-phosphate, but unaffected at higher substrate levels^61^, and we did not observe the *E. coli* GlgC directly affected by ppGpp in our PISA screen.

We further studied how carboxysomes respond to ppGpp *in vivo*. In the ppGpp overproduction strains, carboxysomes aggregated within 24 hours of induction. Although CO₂ fixation was not directly measured, the strong growth arrest suggests reduced carbon metabolism. Mutants of the McdAB carboxysome positioning system in *Syn*7942 have been shown to result in carboxysome aggregation and have been linked to a reduced activity of main carbon fixation enzyme Rubisco under some conditions^51,53^. Yet, the cell localization of the aggregates observed in this study differs from that observed in the McdAB perturbation mutant.

The carboxysome mispositioning we observe during ppGpp overproduction is not observed under conditions of stationary growth, diel light cycles, or phosphate stress^52^, which are conditions ppGpp typically accumulates in (cyano)bacteria. However, the grouping and position of carboxysomes following ppGpp overaccumulation matches that seen in prolonged dark treatment (5 days)^52^. Unlike stationary growth and phosphate stress, prolonged darkness is the only known condition in which the carboxysome positioning protein McdA localizes to the carboxysome in distinct foci. The similar localization pattern of ppGpp induced carboxysome foci and McdA also co-localizing with these aggregates, points to ppGpp playing a key role in dark stress that may affect McdA, leading to co-localization at the carboxysome. Together, this study shows that ppGpp modulates both the stability of specific carboxysome proteins and the spatial organization of carboxysomes *in vivo*, potentially representing a regulatory mechanism linking cellular stress responses to CO₂ fixation machinery.

In addition, we examined the tricarboxylic acid (TCA) cycle and found that ppGpp activates citrate synthase (GltA) in *Syn*6803. Flux through the oxidative TCA cycle is low in cyanobacteria, and their citrate synthase is phylogenetically distinct from heterotrophic counterparts, with unique regulation including inhibition by phosphoenolpyruvate and activation by ADP, MgCl₂, and CaCl₂^48^. We also observed that ppGpp alters citrate synthase oligomerisation. In contrast, *Syn*7942 citrate synthase was not affected by ppGpp in our PISA screen. The citrate synthase in *Syn*7942, but not *Syn*6803, was recently shown to form fractals *in vivo*^62^, and *Syn*7942 citrate synthase activity was previously reported to be unaffected by ppGpp overproduction^9^.

Nitrogen starvation, linked to ppGpp regulation, leads to increased citrate levels in *Syn*6803^63^. Citrate strongly inhibits the oxidative pentose phosphate pathway by targeting the enzymes glucose-6-phosphate dehydrogenase and 6-phosphogluconate dehydrogenase in *Syn*6803, and serves as a metabolic control point to avoid degradation of glycogen reserves during cell dormancy induced by nitrogen limitation^63,64^. The regulation of citrate synthase by ppGpp reported here may provide a connection between nitrogen starvation and citrate-mediated control of carbon flux.

Together, this study provides a new angle on ppGpp-mediated regulation in photosynthetic organisms. By applying a high-throughput chemoproteomic approach, we identified conserved and novel ppGpp targets, extending our understanding of how this secondary messenger coordinates stress responses in photosynthetic systems.

## Methods

### Bacteria and plant cultivation

*A. thaliana* plants were grown in soil at 100 μmol photons m^−2^ s^−1^ in 12 h day/night cycles. Chloroplasts were extracted after 5 weeks using a Percoll gradient to separate intact and broken chloroplasts as previously described ^28^. Chloroplasts were illuminated at 400 µmol·s^−1^·m^−2^ for 10 min before freezing in liquid nitrogen and stored at −80 °C in 50 mM HEPES-KOH (N-(2-hydroxyethyl)piperazine-N′-(2-ethanesulfonic acid) pH 8.03 mM MgSO_4_, 0.3 M sorbitol.

*Syn*6803 and *Syn*7942 were grown at 50 μmol photons m^−2^ s^−1^ in BG-11 media at 1% CO_2_. Cells were harvested between OD_730_ 0.4 to 0.8 by centrifugation and washed twice with lysis buffer (100 mM HEPES-KOH pH 8, 3 mM MgCl_2_, 150 mM KCl) before being resuspended in a small amount of lysis buffer. As with chloroplasts, cyanobacteria were exposed to light, frozen in liquid nitrogen, and stored at −80 °C.

*E. coli* DH5α (K12 derivative) were grown in M9 minimal media at 37 °C and harvested by centrifugation at OD_600_ 0.5, washed twice with lysis buffer, and frozen in liquid nitrogen. Cells were stored at −80 °C until use.

### PISA assay

The PISA assay was performed as previously described with minor modifications^28^. Frozen chloroplast and cyanobacteria aliquots were thawed on ice and NP-40 Surfact-Amps™ Detergent Solution (ThermoFisher) was added to 0.8% v/v. Chloroplasts and cyanobacteria were lysed using a FastPrep-24 5G (45 s, 6.5 m/s) with 3 cycles for chloroplasts and 4 for cyanobacteria. Samples were cooled on ice between cycles to prevent heat. After lysis, cell debris was removed by centrifugation at 21,000 *x* g for 5 min at 4 °C followed by removal of endogenous metabolites using Zeba Spin Desalting Columns (ThermoFisher). After protein quantification using Bradford assay, lysates were split into 200 µg protein replicates in lysis buffer (100 mM HEPES-KOH pH 8.03 mM MgCl_2_, 150 mM KCl), using quadruplicates for each treatment. Samples were incubated with 0, 200, 600, and 1000 μM ppGpp at a protein concentration of 1 µg/µL for 10 min. During the incubation with ppGpp, each replicate was split into 16 aliquots. After incubation, aliquots were heated in a thermocycler between 40-55 °C for chloroplasts and 40-60 °C for cyanobacteria and *E. coli* for 3 min (in total 16 temperature points). Proteins were precipitated at room temperature for 6 min, and then pooled back into the initial sample replicates. Insoluble protein was removed through ultracentrifugation at 150,000 *x* g for 30 min.

Following ultracentrifugation, soluble protein fractions were reduced in 10 mM dithiothreitol (DTT) for 45 min and alkylated in 17 mM iodoacetamide (IAA) for 30 min in darkness. Proteins were digested overnight using Trypsin/Lys-C Protease Mix (Thermo Scientific) at a 1:50 protease:protein ratio. Digestion was quenched by addition of formic acid to pH ∼2.0 before desalting using stage tips packed with six layers of C18 Empore™ SPE Disks (Merck). Desalted peptides were dried in a Speed Vac at 45 °C for approximately 40 min. For TMT labeling, 12.5 µg of each peptide sample was resuspended in 20 µL 20 mM EPPS buffer pH 8.5 and labeled with 0.1 mg TMTpro™ 16plex (Thermo Scientific). Labeled peptides were fractionated into 6 fractions using Pierce™ High pH Reversed-Phase Peptide Fractionation Kit (Thermofisher) and dried in a Speed Vac at 45 °C for approximately 3 h. Samples were stored at −20 °C before LC-MS injection.

### Limited proteolysis assay

Cells of the *Syn*6803 ΔSpoT strain were lysed without detergent and cleared of endogenous metabolites as described in the PISA assay. Lysate (45 μg) and 5 μg recombinant PyrB protein (5 μg) were incubated with and without 500 μM ppGpp at a concentration of 1 μg/μL for 10 min at 25 °C in quadruplicates. Proteinase K (0.5 μg) was added to initiate proteolysis and incubated for an additional 5 min at 25 °C. Proteinase K activity was terminated by heating to 95 °C for 10 min. Proteins were then reduced (10 mM DTT, 45 min), alkylated (17 mM IAA, 30 min darkness), and digested with Trp/LysC mix as previously described. The peptides were desalting using C18 stage tips and dried in a speed vac at 45 °C. Samples were stored at −20 °C before LC-MS injection.

### LC-MS

Peptides were resuspended in 0.1% formic acid and analysed using a Q-Exactive HF Hybrid Quadrupole-Orbitrap Mass Spectrometer with an UltiMate 3000 RSLCnano System with an EASY-Spray ion source. Peptides were loaded on a C18 Acclaim PepMap 100 trap column (75 μm x 2 cm, 3 μm, 100 Å, Thermo Fisher Scientific) at a flow rate of 7 μL per min, with 3% acetonitrile, 0.1% formic acid, and 96.9% water as solvent A. An ES802 EASY-Spray PepMap RSLC C18 Column (75 μm x 25 cm, 2 μm, 100 Å, Thermo Fisher Scientific) with a flow rate of 0.7 μL per min was used to separate the peptides.

PISA assay peptides were separated with a 120 min linear gradient going from 1% to 32% solvent B (95% acetonitrile, 0.1% formic acid, 4.9% water). The peptides were measured using a single full scan (resolution 120,000 at 200 m/z, mass range 350-1500 m/z) followed by 15 MS2 DDA scans using the 15 most abundant peptides, with dynamic exclusion set to 30 s. For MS2 scans, a resolution of 60,000 at 200 m/z was maintained, with an isolation window of 0.7 m/z. Precursor ions were fragmented using high-energy collision-induced dissociation at an NCE of 30. Maximum injection time and automatic gain control were set to 50 ms and 3E6 for MS1, and 120 ms and 1E5 for MS2, respectively.

Limited proteolysis peptides were separated using a 40 min linear gradient from 1% to 32% elution solvent. Peptides were measured using one full scan (resolution 30,000 at 200 m/z, mass range 300–1200 m/z) followed by 30 MS2 DIA scans (resolution 30,000 at 200 m/z, mass range 350–1000 m/z) with an isolation window of 10 m/z. Precursor ion fragmentation was performed with high-energy collision-induced dissociation at an NCE of 26. Maximum injection times for the MS1 and MS2 were 105 ms and 55 ms, and automatic gain control was set to 3E6 and 1E6, respectively.

### Data analysis

For protein identification the following UniProt proteomes were used as reference libraries: UP000006548 (*A. thaliana*, limited to reviewed entries), UP000001425 (*Syn*6803), UP000002717 (*Syn*7942), and UP000000625 (*E. coli* strain K12).

For PISA samples, protein identification and quantification were conducted using MaxQuant version 2.2.0.0. Variable modifications were set as oxidation of methionine and N-terminus acetylation, while carbamidomethylation of cysteine was set as a fixed modification. Up to two missed cleavages were permitted and a false discovery rate of 1% was applied.

For limited proteolysis samples the data was searched using the EncyclopeDIA version 1.2.2 search engine. The prosit intensity prediction model “Prosit_2020_intensity_hcd” was used to generate a predicted peptide library for *Syn*6803 from the UniProt proteome with the *A. thaliana* PyrB sequence added. A maximum of two missed cleavages and a false discovery rate of 1% was used.

Proteins were median normalized and fold-changes were calculated using MSstatsTMT (v. 2.4.1) and MSstats (v. 4.4.1) packages in R^65^. The p-values were adjusted for multiple hypothesis testing using the Benjamini-Hochberg method, with an adjusted p-value threshold of 0.05. Proteins detected in fewer than 3 replicates were excluded from statistical analysis.

### Recombinant protein expression and purification

Proteins were codon-optimized for *E. coli* and synthesized as geneblocks (Integrated DNA Technologies, codon-optimised for *E. coli*) or amplified from genomic DNA and cloned into a pET expression vector with a 6xHis-tag or strep-tag. Transit peptides were excluded from *A. thaliana* genes based on Uniprot sequence annotation. *Syn*6803-CcaA was truncated by 51 residues at the C-terminus and N-terminally fused to a 6xHis-MBP domain to increase solubility. *Syn*6803-GlgC was N-terminally truncated by 10 residues as the UniProt sequence is likely misannotated^66^. Plasmids were transformed into *E. coli* BL21-DE3 or BL21-AI after verification by sequencing. Expression details and protein sequences are available in Supplementary Table 1.

BL21 cells were cultured in low salt LB media at 37 °C until OD_600_ 0.3-0.8. Expression was induced with 0.5 mM IPTG (BL21-DE3) or 0.1 % arabinose (BL21-AI) and grown at 18-20 °C overnight. Cultures were harvested by centrifugation and stored at −20°C until use. BL21-DE3 cells were lysed using B-PER Complete Bacterial Protein Extraction Reagent (Thermo Fisher Scientific). BL21-AI cells were resuspended in protein purification binding buffer (see below) supplemented with 0.1 mg/mL lysozyme, 0.1 μg/mL benzonase (Merck) and 1 mM phenylmethylsulfonyl fluoride (PMSF) and lysed by sonication at 65% amplitude with two rounds of 10 s pulse 15 s pause for 1 min (VibraCell, Sonics). Lysates were cleared by centrifugation.

For *A. thaliana* PyrB, *A. thaliana* Amy3, *Syn*6803 GlnB, *Syn*6803 GlgC, *Syn*6803 PyrE, *Syn*6803 GltA, and *Syn*7942 CcmL, the lysed cells were filtered through a 0.2 μm filter and purified using an ÄKTA start™ protein purification system equipped with a 1 mL HisTrap Fast Flow column (Cytiva). The column was washed with 15 column volumes of wash buffer (50 mM Tris-HCl, 500 mM NaCl, 20 mM imidazole, pH 8.0), eluted with a stepwise gradient of elution buffer (50 mM Tris-HCl, 500 mM NaCl, 300 mM imidazole, pH 8.0) and fractions containing the target protein were collected.

*Syn*6803-CcmK1, *Syn*6803-CcmK2, *Syn*6803-CcmP and *Syn*6803-CcmL were batch purified via His-tag affinity using Ni Sepharose 6 Fast Flow (Cytiva) resin. The resin was pre-equilibrated with binding buffer (50 mM Tris-HCl, 300 mM NaCl, 20 mM imidazole, pH 8.0) and incubated with the clarified lysate for 1 h on ice. Nonspecific proteins were washed away with 4×50 mL of wash buffer (50 mM Tris-HCl, 300 mM NaCl, 60 mM imidazole pH 8.0) by pelleting the resin (centrifugation 500 x *g*, 5 min) and then resuspending it in fresh wash buffer. Proteins were eluted with 10 mL elution buffer (50 mM Tris, 300 mM NaCl, 300 mM imidazole, pH 8.0).

*Syn*6803-CcaA and *Syn*6803-CcmK4 were purified via His-tag affinity using a 5 mL HisTrap Fast Flow column (Cytiva) attached to a syringe pump (AL-4000, KF technologies) at a flow rate of 5 mL/min. Clarified lysate was loaded to the column, pre-equilibrated with His-binding buffer and nonspecifically bound proteins were washed away with 60 mL of His-wash buffer. Bound proteins were thereafter eluted with 30 mL of His-elution buffer. *Syn*6803-CcmN was purified via strep-affinity using a 5 mL StrepTrap HP column (Cytiva) attached to a syringe pump at a flow rate of 5 mL/min. Clarified lysate was applied to the column, pre-equilibrated with strep-binding buffer (50 mM Tris-HCl, 300mM NaCl pH 8.0). The column was thereafter washed with 60 mL strep-binding buffer and bound proteins were eluted with 30 mL strep-elution buffer (50 mM Tris-HCl, 300 mL NaCl, 2.5 mM desthiobiotin, pH 8.0).

Following purification, proteins were concentrated using 10 or 30 kDa Mw cut-off centrifugal filters (Millipore) and buffer exchanged using PD-10 desalting column (Cytiva) into storage buffer. Syn6803 carboxysomal proteins (CcmL, CcmP, CcmL, CcmK1, CcmK2, CcmK4 and CcaA) were stored in phosphate buffered saline (PBS) buffer pH 7.4 with 10% v/v glycerol and all other proteins were stored in 50 mM Tris-HCl buffer pH 8.0. Proteins were flash frozen in liquid nitrogen and stored at −80 °C in single-use aliquots. Purity was assessed by SDS-PAGE and protein concentrations were determined by A_280_ using a NanoDrop2000 (Thermo Scientific) and the theoretically calculated extinction coefficient (ProtParam).

### In vitro measurement of thermal stability

The *in vitro* thermal stability of purified proteins was measured using a Nano Differential Scanning Fluorimetry (F350/F330) using a Prometheus NT.48 (NanoTemper). Proteins were pre-incubated for 10 min at room temperature at the tested conditions using 0.2-0.5 mg/mL protein in 50 mM Tris-HCl pH 8.0 unless indicated otherwise in figure texts. The excitation power was adjusted to achieve a fluorescence between 5,000-15,000 counts. Protein stability was continuously measured via 330 and 350 nm fluorescence and scattering, with heating from 25 to 100 °C at an increase of 1 °C per min.

### Kinetic assays of plant Aspartate carbamoyltransferase

Activity of aspartate carbamoyltransferase (*A. thaliana* PyrB) was measured colorimetrically as described previously^67^. The assays were conducted in a total volume of 60 μL in 50 mM Tris-Acetate buffer at pH 8.3 with a final *PyrB* concentration of 1 ng/μL (26 nM). The enzyme was pre-incubated with variable aspartate concentrations at 25 °C and ppGpp or GDP for 5 min before the reaction was initiated by addition of carbamoyl phosphate to a final concentration of 5 mM. After 5 min, the reactions were quenched with an equal volume color reagent (5 g/L antipyrine in 50 % sulphuric acid and 8 g/L 2,3-butadionemonoxime, mixed at a 2:1 ratio) and were placed in darkness for 16 h. Subsequently, the reactions were heated to 45 °C for 20 min under normal lab ceiling illumination prior to absorbance measurements at 466 nM in a plate reader.

### Molecular docking

The binding of ppGpp to *A. thaliana* PyrB was simulated by DynamicBind model version 2 using the Neurosnap online platform ^68^ using the crystal structure bound to UMP and the apoform (PDB structure 6YPO and 6YY1 respectively). The best prediction out of 10 models was selected based on cLDDT score, using 20 inference steps and allowing for post-prediction relaxation and noise during reverse diffusion.

### Assay of Glucose-1-phosphate adenylyltransferase

Glucose-1-phosphate adenylyltransferase was assayed by measuring formation of ADP-glucose from glucose-1-phosphate (G1P) and ATP. In *A. thaliana*, plants were grown as described above for 4 weeks. Leaves were harvested in the middle of the day cycle, and immediately frozen in liquid nitrogen and grinded to a powder on dry ice using mortar and pestle. Grinded leaves (100 mg) were resuspended in 1 mL 100 mM ammonium bicarbonate buffer pH 7.5 with 5 mM MgCl_2_ and 20 mM DTT, and incubated on ice for 30 min. The leaf lysate was cleared by centrifugation at 21,000 *x* g for 5 min and the supernatant was collected. The ADP-glucose formation was assayed at 2 mM G1P, and ATP (0-2 mM) by adding 25 μL leaf extract (0.4 μg/μL protein) into 25 μL solution containing 2X concentrated ATP, G1P, and ppGpp or milliQ in triplicates. The solution was incubated at 21 °C for 10 min, and quenched by boiling for 3 min. Precipitates were removed by centrifugation and the supernatant was stored at 4 °C until HPLC injection the same day.

ADP-glucose formation in *Syn*6803 was assayed in the same way using recombinant GlgC at a final concentration of 25 ng/μL (0.5 μM) protein in 0.125-1 mM G1P and 1 mM ATP in triplicates in HEPES-KOH buffer at pH 8.0 with 5 mM MgCl_2_. The reaction was incubated at 30 °C for 10 min and quenched by boiling.

ADP-glucose was detected on a Vanquish UHPLC system (Thermo Scientific) equipped with a Acclaim Vanquish C18 column (150 x 2.1 mm, 2.2 μm particle size) and UV-detector monitoring 260 nm. Nucleotides were separated using a 27 min isocratic gradient with 31 mM triethylammonium bicarbonate, 30 mM ammonium bicarbonate buffer pH 7.5 at a flow rate of 0.6 mL/min, injecting 20-25 μL sample. The buffer was prepared the same day of injection and stored on ice. Between each sample, the column was flushed for 3 min with 75% acetonitrile and equilibrated for 5 min. Samples from the highest and lowest ATP concentration was injected using a 65 min isocratic gradient to verify that ATP was not depleted during the assay in lysates and standards of ATP, ADP, ADP-glucose and AMP were used to verify separation of nucleotides (Supplementary Figure 4B).

### Isothermal titration calorimetry

Experiments were performed in an iTC-200 calorimeter (MicroCal, Malvern-Panalytical) at 25 °C. The reaction cell contained 880 ng/μL (60 μM) *Syn*6803 PII protein (GlnB) and the titration syringe contained 1 mM ppGpp or ATP. Both protein and titrant were in 10 mM Tris buffer pH 8.0 with 5 mM MgCl_2_. Titration experiments consisted of 19 injections of 2 µl each, with an initial injection of 0.4 μL titrant. Data analysis was performed with Origin 7 (OriginLab) using non-linear least-squares regression analysis to estimate dissociation constants. Titration of 1 mM ppGpp or ATP into the buffer without protein was used to normalise the signal for each respective experiment.

### Kinetic assays of citrate synthase

*Syn*6803 citrate synthase (GltA) activity was assayed by measuring the formation of CoA from the condensation of oxaloacetate and acetyl-CoA. The reaction mixture contained 3.8 ng/μL (80 nM) *Syn*6803 GltA, 1 mM oxaloacetate, 5 mM MgCl₂, 150 mM KCl, and 50 mM Tris-HCl (pH 8.0), with or without 220 μM ppGpp (final assay concentration of 200 μM), and pre-incubated for 10 min at room temperature. DTNB (5,5′-dithiobis-(2-nitrobenzoic acid)) was added to 200 μM to react with the free thiol group of CoA, producing a yellow-colored product. The reaction was initiated by addition of acetyl-CoA (0–200 μM) to a final volume of 100 μL, and initial rates were measured continuously at 412 nm in quadruplicate.

### Mass photometry

*Syn*6803 GltA oligomerization was measured using a TwoMP mass photometer (Refeyn). Sample well cassettes were set up on a microscopic cover slip mounted on the mass photometer using immersion oil. To focus the instrument with droplet dilution, 19 μL 50 mM Tris-HCl buffer pH 8.0 with 5 mM MgCl2 was added to the wells with and without 250 μM ppGpp. Then 1 µL of 1 µM purified protein was added and mixed thoroughly. Oligomeric state populations were recorded using AcquireMP with a 1-minute acquisition time and analyzed with DiscoverMP by quantifying protein binding events and Gaussian fitting. Molecular mass was determined by comparing histogram contrast values to the calibration standard measured on the same day, using bovine serum albumin and thyroglobulin.

### Assay of alpha-amylase

*A. thaliana* alpha-amylase (Amy3) activity was assayed using EnzChek™ *Ultra* Amylase Assay Kit (Thermo Fisher) which releases fluorescent DQ starch molecules upon amylase digestion. Recombinant Amy3 (30 ng/μL) were added in quadruplicates to DQ starch substrate solution (0-50 μg/mL) with and without ppGpp in 50 mM Tris buffer pH 7.5. Amy3 was reduced on ice for 1 h before the assay using 20 mM DTT. Amylase activity was then measured continuously at an excitation of 495 nm and emission at 520 nm for 30 minutes.

### Cloning of YjbM *Syn*7942 overexpression strain

The gene YjbM was constructed using Gibson Assembly into a plasmid under a Ptrc promoter with a theophylline inducible riboswitch and transformed into neutral site II of a *Syn*7942 strain expressing RbcS tagged with monomeric Turquoise2 (RbcS-mTurquoise2) under the native RbcS promoter in neutral site I^51,69^.

### Culture conditions YjbM *Syn*7942 overexpression strain

Cultures of the *Syn*7942 YjbM strain were grown in a BG-11 medium (Sigma-Aldrich, C3061) buffered with 1 g/L HEPES at either pH 8.3 or 6.3 with NaOH, as indicated in the text. Cells were grown at ∼125 μmol photons m^−2^ s^−1,^ supplemented with 2% CO_2_ at 30 °C. To increase consistency in between-day experiments, cultures were maintained with daily back dilution to an OD_750_ of 0.3.

### Measurement of pigments

Chlorophyll A (ChlA) and carotenoid content was measured in triplicates in *Syn*7942 cultures with an OD_750_ between 0.3-1. Cells (1 mL) were harvested by centrifugation, resuspended in 1 mL ice-cold methanol and stored for 30 min in darkness at 4 °C. The absorbance was measured at 470, 665, and 720 nm. ChlA and carotenoid content were calculated as below and normalised to OD_750_. Absorbance at 720 nm was used to correct for background absorbance.

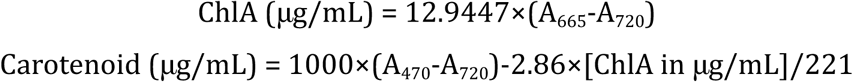

Phycocyanin content was determined in triplicate measurements using 1 mL cultures harvested by centrifugation above. Cells were lysed in 200 μL phosphate buffered saline pH 7.4 (PBS) via one cycle of bead beating as described earlier. The volume was then adjusted to 1 mL by addition of PBS, vortexed, and stored on ice in darkness for 1 h. The lysates were cleared by centrifugation at 17,000 *x* g for 5 minutes at 4 °C. The absorbance was measured at 615, 652 and 720 nm, and a full spectra measurement in the range of 550-670 nm. Phycocyanin content was calculated as below^70^.

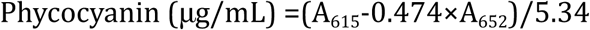

### Fluorescence microscopy and image analysis

Fluorescence images were taken with a Zeiss Axio Observer D1 microscope (63×1.3NA) with an Axiocam 503 (mono-chrome) camera using light from X-Cite 120Q (Lumen Dynamics, Mississauga, Canada). Chla fluorescent signal was taken using filter set 43 (000000-1114-101): excitation BP 545/25, emission BP 605/70, and beam splitter FT570. The turquoise fluorescent signal was taken using filter set 47 (000000-1196-682): excitation BP 436/20, emission BP 480/40, and beam splitter FT455. Images were processed in Fiji using background subtraction 20-50 with a sliding paraboloid^71^. These data were further analyzed using MicrobeJ 5.13p^72^. In each cell line, cell length detection was performed using the rod-shaped descriptor and thresholding set to 0.4 μm < area < 7, 2 µm < length, 0.6 μm < width range < 2 μm, 0 μm < width variation < 0.2 μm, and 0.15 μm < circularity amplitude. Carboxysome detection was performed using the Maxima point function with a tolerance of 200, a Z-score of 50, and an intensity minimum of 700. Associations such as inside, location, distance, and spacing were all used at the default setting. Graphs and statistical analyses were generated with GraphPad Prism 10.4.1.

### Sytox blue dead cell staining

1 mL of *Syn*7942 cells expressing RbcS-mTQ and YjbM with or without theophylline induction were harvested after 24 hours at 0.4 OD750. For dead cell control, 1 mL of WT cells was incubated at 95 °C for 10 min. SYTOX Blue nucleic acid stain was added to the cell suspension to a concentration of 1 µM and incubated in the dark for 40 min. Cells were imaged using a Zeiss Axio Observer D1 microscope as described above and average CFP channel intensity per cell was plotted. Dead cells were identified based on SYTOX blue fluorescence, whereas live cells remained unstained.

### Transmission Electron microscopy

For primary fixation 2 mL of later stage cultures (OD750 ∼0.8) was pelleted and resuspended in 1 mL Glutaraldehyde 2%, paraformaldehyde 2% in PBS pH 7.4. After primary fixation, samples were washed with 0.1M phosphate buffer and post-fixed with 1% osmium tetroxide in 0.1M phosphate buffer, in block stain with 1% uranyl acetate, dehydrated in a gradient series of acetone and infiltrated and embedded in Spurr resin. 70 nm thin sections were obtained with a Enuity Ultramicrotome (Leica Mycrosystems) and post stained with uranyl acetate and lead citrate. Images were taken with JEOL 1400Flash Transmission Electron Microscope (Japan Electron Optics Laboratory, Japan) at an accelerating voltage of 100kV.

## Data and code availability

Code for analysis of PISA data has been deposited to GitHub (https://github.com/emilsporre/pisa (03/25/2025)).

## Acknowledgements

We are grateful to Karen Schriever at the SciLifeLab Drug Discovery & Development facility for access and assistance of isothermal calorimetry measurements. We thank the undergrad students Hing Pan Ng from Uppsala University and Agnes Benjaminsson, Alva Lundmark Brånstad, Elissar Ghanem, Elsa Hesslow from KTH Royal Institute of Technology for assistance with cloning and purification of proteins, and internship student Ann-Christin Branscheid from Technische Universität Berlin for optimization of the limited proteolysis workflow.

## Author contributions

AK, EmS, ÅS, and EPH conceptualized and designed the project. AK performed most of the experiments and wrote the majority of the article together with EPH. Proteomic data analysis script and PyrB assay was carried out by EmS. Carboxysomal strain cloning, microscopy, cell quantification, and pigment measurement was carried out by RR. Proteins were cloned and purified by AK, EE, and NV. Citrate synthase cloning and kinetic assays were set up by JLR and AK. CG carried out mass photometry analysis. FE, ES, CB, ÅS, DD and EPH provided supervision. All authors have reviewed and approved of the final manuscript.

