## Supplementary Figures and Tables for "Identifying Novel Targets of the Stringent Response in Plants and Cyanobacteria using chemoproteomics"

**Supplementary Table 1.** Protein sequences and expression details for recombinant proteins in this study.

| Protein | Origin | Expression | Purification | FASTA sequence |
|---|---|---|---|---|
| *A. thaliana*<br>PyrB | Geneblock | pET28a(+)<br>BL21-DE3 | N-terminal<br>6xHis-tag | CHAMQAGTRELKKFELSDVIEGKQFDREMLSAIFDVARE<br>MEKIEKSSSQSEILKGYLMATLFYEPSTRTRLSFESAMKRL<br>GGEVLTTENAREFSSAAKGETLEDTIRTVEGYSDIIVMRH<br>FESGAARKAAATANIPVINAGDGPGEHPTQALLDVYTIQS<br>EIGKLDGISVALVGDLANGRTVRSLAYLLAKFKDVKIYFVS<br>PEIVKMKDDIKDYLTSSGVEWEESSDLMEVASKCDVVYQ<br>TRIQRERFGERLDLYEAARGKYIVDKDLLGVMQKKAIIMH<br>PLPRLDEITADVDADPRAAYFRQAKNGLFIRMALLKLLLV<br>GW |
| *A. thaliana*<br>Amy3 | Geneblock | pET28a(+)<br>BL21-DE3 | N-terminal<br>6xHis-tag | SMNKSPVAIRATSSDTAVVETAQSDDVIFKEIFPVQRIEKA<br>EGKIYVRLKEVKEKNWELSVGCSIPGKWILHWGVSYVGD<br>TGSEWDQPPEDMRPPGSIAIKDYAIETPLKKLSEGDSFFE<br>VAINLNLESSVAALNFVLKDEETGAWYQHKGRDFKVPLV<br>DDVPDNGNLIGAKKGFGALGQLSNIPLKQDKSSAETDSIE<br>ERKGLQEFYEEMPISKRVADDNSVSVTARKCPETSKNIVSI<br>ETDLPGDVTVHWGVCKNGTKKWEIPSEPYPEETSLFKNK<br>ALRTRLQRKDDGNGSFGLFSLDGKLEGLCFVLKLNENTW<br>LNYRGEDFYVPFLTSSSSPVETEAAQVSKPKRKTDKEVSA<br>SGFTKEIITEIRNLAIDISSHKNQKTNVKEVQENILQEIEKL<br>AAEAYSIFRSTTPAFSEEGVLEAEADKPDIKISSGTGSGFEI<br>LCQGFNWESNKSGRWYLELQEKADELASLGFTVLWLPP<br>PTESVSPEGYMPKDLYNLNSRYGTIDELKDTVKKFHKVGI<br>KVLGDAVLNHRCAHFKNQNGVWNLFGGRLNWDDRAVV<br>ADDPHFQGRGNKSSGDNFHAAPNIDHSQDFVRKDIKEW<br>LCWMMEEVGYDGWRLDFVRGFWGGYVKDYMDASKPYF<br>AVGEYWDSLSYTYGEMDYNQDAHRQRIVDWINATSGAA<br>GAFDVTTKGILHTALQKCEYWRLSDPKGKPPGVVGWWP<br>SRAVTFIENHDTGSTQGHWRFPEGKEMQGYAYILTHPGT<br>PAVFFDHIFSDYHSEIAALLSLRNRQKLHCRSEVNIDKSER<br>DVYAAIIDEKVAMKIGPGHYEPPNGSQNWSVAVEGRDYK<br>VWETS |
| *Syn*6803<br>GlnB | Geneblock | pET28a(+)<br>BL21-DE3 | N-terminal<br>6xHis-tag | MKKVEAIIRPFKLDEVKIALVNAGIVGMTVSEVRGFGRQK<br>GQTERYRGSEYTVEFLQKLKIEIVVDEGQVDMVVDKLVSA<br>ARTGEIGDGKIFISPVDSVVRIRTGEKDTEAI |

| | | | | |
|---|---|---|---|---|
| *Syn*6803 GlgC* | Genomic DNA | pET28a(+) BL21-DE3 | N-terminal 6xHis-tag | VKRVLAIILGGGAGTRLYPLTKLRAKPAVPLAGKYRLIDIP VSNCINSEIVKIYVLTQFNSASLNRHISRAYNFSGFQEGFVE VLAAQQTKDNPDWFQGTADAVRQYLWLFREWDVDEYL ILSGDHLYRMDYAQFVKRHRETNADITLSVVPVDDRKAP ELGLMKIDAQGRITDFSEKPQGEALRAMQVDTSVLGLSAE KAKLNPYIASMGIYVFKKEVLHNLLEKYEGATDFGKEIIPD SASDHNLQAYLFDDYWEDIGTIEAFYEANLALTKQPSPDF SFYNEKAPIYTRGRYLPPTKMLNSTVTESMIGEGCMIKQC RIHHSVLGIRSRIESDCTIEDTLVMGNDFYESSSERDTLKA RGEIAAGIGSGTTIRRAIIDKNARIGKNVMIVNKENVQEAN REELGFYIRNGIVVVIKNVTIADGTVI |
| *Syn*6803 PyrE | Genomic DNA | pET28a(+) BL21-DE3 | N-terminal 6xHis-tag | MTDLTLAALKTAPLAQVRQYLLHLLATHAYKEGDFILSSG QPSTYYINGKLVTLRAEGALAIGRLLLTELPDQVEAVAGL TLGADPIVSAVSTVSAYEEKPVVALIIRKEAKGHGTKAYIE GPELAPGTKVVVLEDVVTTGKSAMLAVERLRNAGYQVDT VISLVDRQQGGREFYQSQGLTFQALFTIGDIQQVYRQK |
| *Syn*6803 GltA | Genomic DNA | pET28a(+) BL21-DE3 | N-terminal 6xHis-tag | MMTDNEVFKEGLAGVPAAKSRVSHVDGTDGILEYRGIRIE ELAKSSSFIEVAYLLIWGKLPTQAEIEEFEYEIRTHRRIKYH IRDMMKCFPETGHPMDALQTSAAALGLFYARRALDDPK YIRAAVVRLLAKIPTMVAAFHMIREGNDPIQPNDKLDYAS NFLYMLTEKEPDPFAAKVFDVCLTLHAEHTMNASTFSAR VTASTLTDPYAVVASAVGTLAGPLHGGANEEVLNMLEEI GSVENVRPYVEKCLANKQRIMGFGHRVYKVKDPRAIILQD LAEQLFAKMGHDEYYEIAVELEKVVEEYVGQKGIYPNVDF YSGLVYRKLDIPADLFTPLFAIARVAGWLAHWKEQLSVN KIYRPTQIYIGDHNLSYVPMTERVVSVARNEDPNAII |
| *Syn*6803 CcmL | Genomic DNA | pET14b BL21-AI | C-terminal 6xHis-tag | MQLAKVLGTVVSTSKTPNLTGVKLLLVQFLDTKGQPLER YEVAGDVVGAGLNEWVLVARGSAARKERGNGDRPLDAM VVGIIDTVNVASGSLYNKRDDGR |
| *Syn*6803 CcmN | Genomic DNA | pET14b BL21-AI | N-terminal strep-tag | MQLPPVHSVSLSEYFVSGNVIIHETAVIAPGVILEAAPDCQI TIEAGVCIGLGSVISAHAGDVKIQEQTAIAPGCLVIGPVTIG ATACLGSRSTVFQQDIDAQVLIPPGSLLMNRVADVQTVGA SSPTTDSVTEKKSPSTANPIAPIPSPWDNEPPAKGTDSPS DQAKESIARQSRPSTAEAAEQISSNRSPGESTPTAPTVVTT APLVSEEVQEKPPVVGQVYINQLLLTLFPERRYFSS |
| *Syn*6803 CcmP | Genomic DNA | pET14b BL21-AI | C-terminal 6xHis-tag | MGIELRSYVYLDSLQSQHAAYIGTVASGFLPLPGDCSLWV EVSPGIEINRITDIALKAAVVRPGVLFVERLYGLLEIHASNQ |

| | | | | |
|---|---|---|---|---|
| | | | | GEVRAAGQAILAYIGAKASDCIKPKVVSSQIIRNIDAYQTQL INRNRRGHMLLAGQTLFVLEVQPAAYASLAANEAEKSASI NILQVSSIGSFGRLYLGGEERDIKAGARAAIAAIENAPGKV PTLEGKNE |
| *Syn*6803 CcmK1 | Genomic DNA | pET14b BL21-AI | N-terminal 6xHis-tag | MSIAVGMIETLGFPAVVEAADSMVKAARVTLVGYEKIGSG RVTVIVRGDVSEVQASVTAGIENIRRVNGGEVLSNHIIARP HENLEYVLPIRYTEAVEQFREIVNPSIIRR |
| *Syn*6803 CcmK2 | Genomic DNA | pET14b BL21-AI | N-terminal 6xHis-tag | MSIAVGMIETRGFPAVVEAADSMVKAARVTLVGYEKIGSG RVTVIVRGDVSEVQASVSAGIEAANRVNGGEVLSTHIIARP HENLEYVLPIRYTEEVEQFRTY |
| *Syn*6803 CcmK4 | Genomic DNA | pET14b BL21-AI | N-terminal 6xHis-tag | MSAQSAVGSIETIGFPGILAAADAMVKAGRITIVGYIRAGS ARFTLNIRGDVQEVKTAMAAGIDAINRTEGADVKTWVIIP RPHENVVAVLPIDFSPEVEPFREAAEGLNRR |
| *Syn*6803 CcaA** | Genomic DNA | pET14b BL21-AI | N-terminal 6xHis-MPB | MQRLIEGLQKFREGYFSSHRDLFEQLSHGQHPRILFICCSD SRVDPNLITQSEVGDLFVIRNAGNIIPPYGAANGGEGAAM EYALVALEINQIIVCGHSHCGAMKGLLKLNSLQEKLPLVY DWLKHTEATRRLVLDNYSHLEGEDLIEVAVAENILTQLK NLQTYPAIHSRLHRGDLSLHGWIYRIEEGEVLAYDGVLHD FVAPQSRINALEPEDEYALHPNS |
| *Syn*7942 CcmL | Genomic DNA | pET28a(+) BL21-DE3 | N-terminal 6xHis-tag | MRIAKVRGTVVSTYKEPSLQGVKFLVVQFLDEAGQALQEY EVAADMVGAGVDEWVLISRGSQARHVRDCQERPVDAAVI AIIDTVNVENRSVYDKREHS |

* N-terminal truncation

** C-terminal truncation

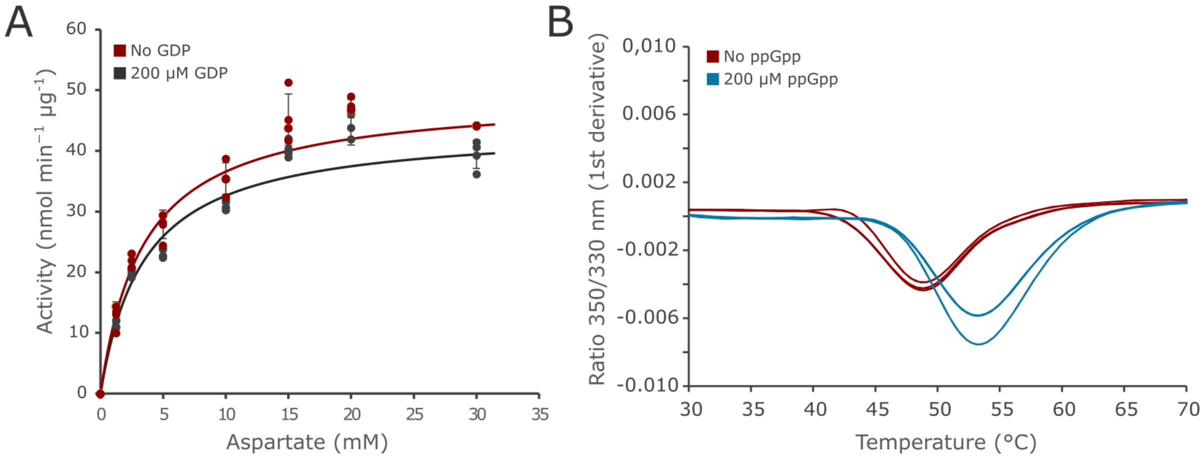


**Supplementary Figure 1**. A) Activity assay of *A. thaliana* PyrB with and without addition of 200 µM GDP, fitted to a Michaelis-Menten curve. Lines represent the mean and standard deviation (SD) of four replicates. B) ppGpp-induced change in thermal stability of Syn6803 PyrE (triplicate measurements) measured by NanoDSF.

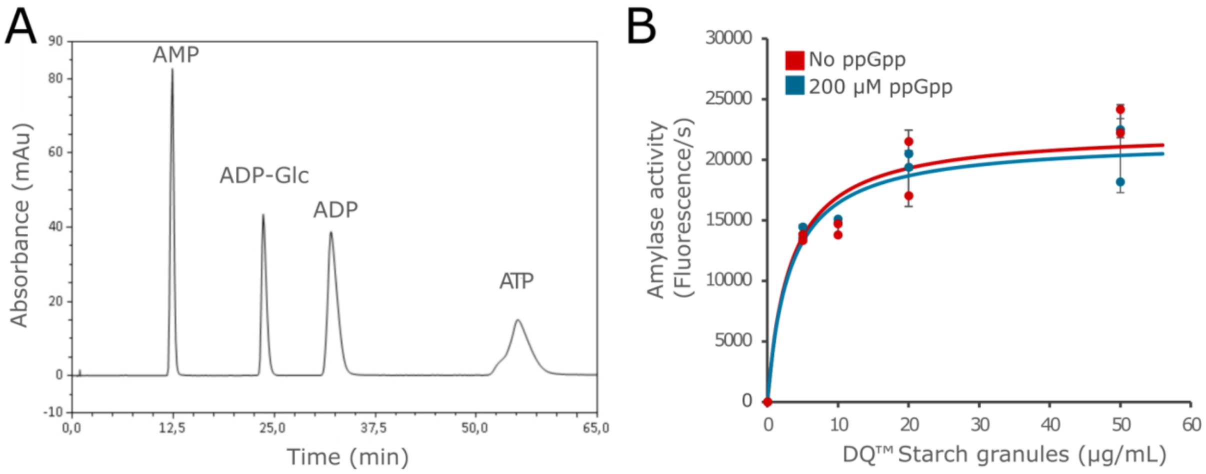


**Supplementary Figure 2**. A) UV-HPLC chromatogram showing detection of AMP, ADP, ADP-glucose, and ATP (100 µM each) used for assaying glucose-1-phosphate adenylyltransferase activity. B) Starch degradation assay with recombinant alpha-amylase 3 (*A. thaliana* Amy3), measuring release of fluorescent dye from DQ™ starch granules in duplicates, fitted to a Michaelis-Menten curve via non-linear regression. Lines represent the mean and SD of two replicates.

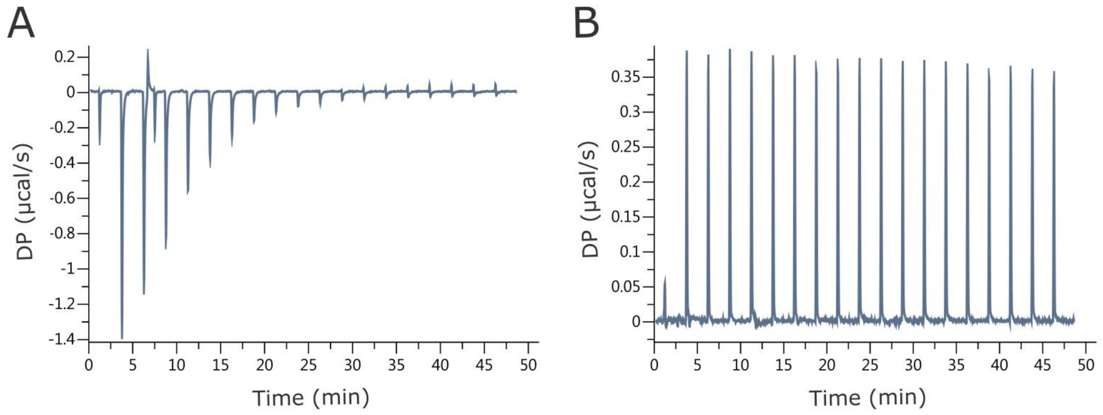


**Supplementary Figure 3.** Isothermal titration calorimetry measurement of 60 µM PII protein (*Syn*6803 GlnB), showing differential power (DP) for titration with A) 1 mM ATP (positive control) and B) 1 mM ppGpp as titrant.

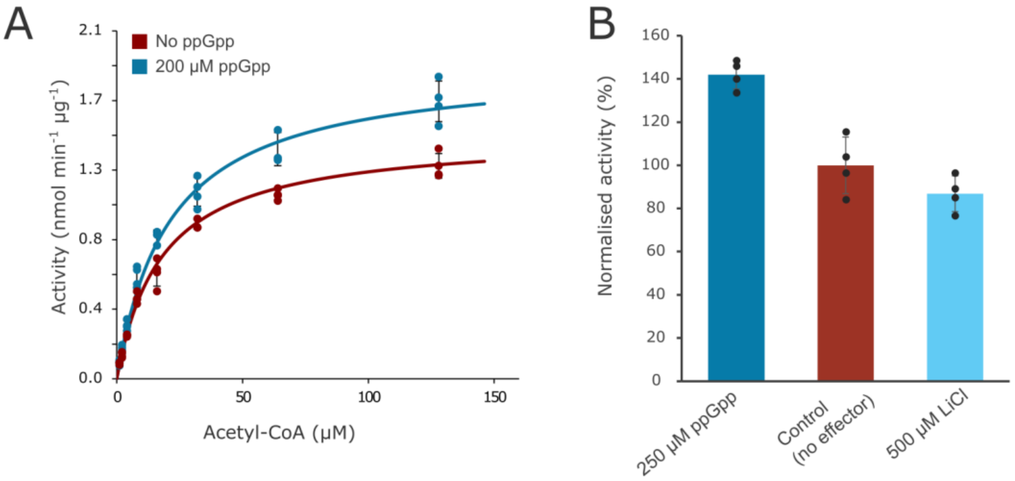


**Supplementary Figure 4.** Kinetic assay of recombinant *Syn*6803 citrate synthase (GltA) without $MgCl_2$ in the reaction buffer. A) Activity in presence of ppGpp without $MgCl_2$ fitted to a Michaelis-Menten curve by non-linear regression. Lines represent mean and SD of four replicates. B) GltA reaction rate at 1 mM oxaloacetate, 150 µM acetyl-CoA, and 3 mM $MgCl_2$, in the presence of ppGpp and LiCl, with activity normalised to the activity without ppGpp and LiCl. Lines represent mean and SD of four replicates.

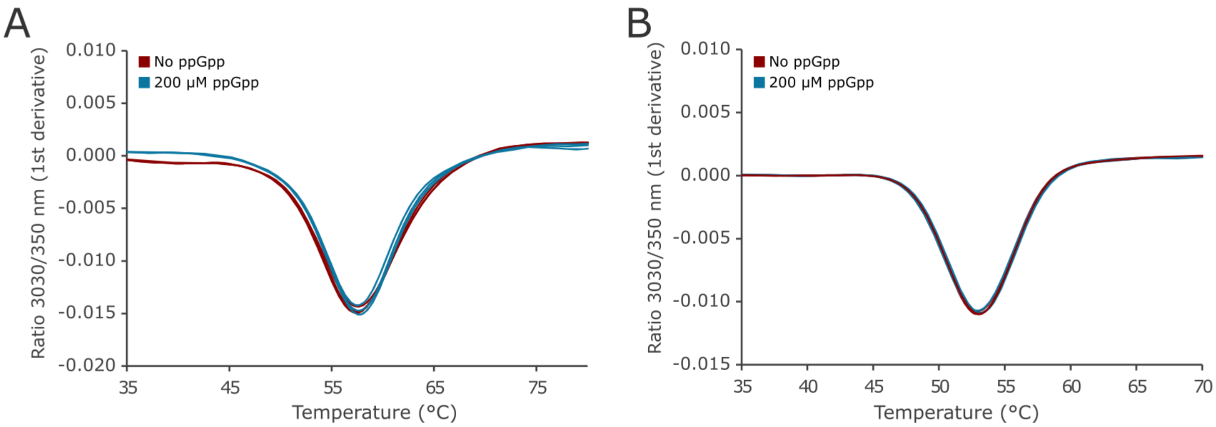


**Supplementary Figure 5.** NanoDSF melting curves of A) Syn7942 CcmL and B) *Syn*6803 CcaA in triplicate replicate measurements.

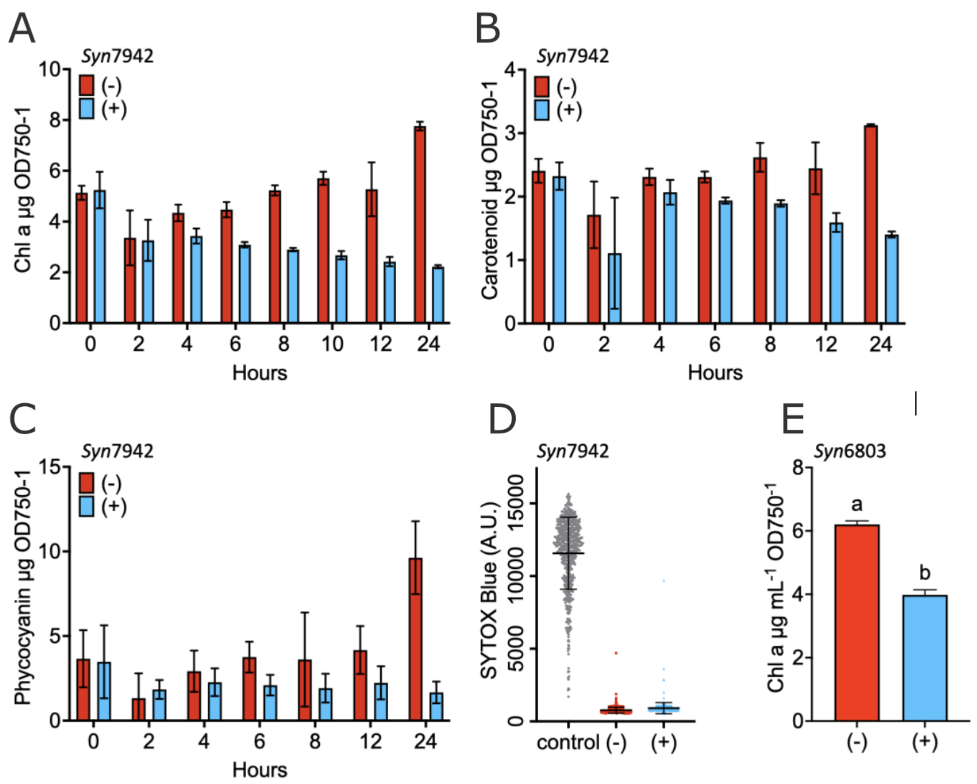


**Supplementary Figure 6.** Pigment content of *Syn*7942 expressing YjbM (RelQ small ppGpp synthase) with (+) and without (-) theophylline as inducer. Pigments were measured under 24 hours after induction (A-C). D) Cell viability analysis of *Syn*7942 via SYTOX blue after 24 hours with (+) or without (-) induction of YjbM. A positive control (grey; heat-inactivated cells) is included to represent expected staining for dead cells. E) Chlorophyll content of *Syn*6803 expressing YjbM (RelQ small ppGpp synthase) with (+) and without (-) theophylline as inducer. Chlorophyll was measured at 48 hours after induction. Bars and errors represent the mean and SD of three or more replicates.
